# Uncovering nitroxoline activity spectrum, mode of action and resistance across Gram-negative bacteria

**DOI:** 10.1101/2024.06.04.597298

**Authors:** Elisabetta Cacace, Manuela Tietgen, Meike Steinhauer, André Mateus, Tilman G. Schultze, Marina Eckermann, Marco Galardini, Vallo Varik, Alexandra Koumoutsi, Jordan J. Parzeller, Federico Corona, Askarbek Orakov, Michael Knopp, Amber Brauer-Nikonow, Peer Bork, Celia V. Romao, Michael Zimmermann, Peter Cloetens, Mikhail M. Savitski, Athanasios Typas, Stephan Göttig

**Author notes:** These authors contributed equally (Elisabetta Cacace, Manuela Tietgen).

## Abstract

Nitroxoline is a bacteriostatic quinoline antibiotic, considered a metal chelator inhibiting the activity of RNA-polymerase^1^. Its clinical indications are limited to uncomplicated urinary tract infections (UTIs), with a clinical susceptibility breakpoint only available for *Escherichia coli*^2^. By testing > 1,000 clinical isolates, here we demonstrate a much broader activity spectrum and species-specific bactericidal activity, including multidrug-resistant Gram-negative bacteria for which therapeutic options are limited due to resistance. By combining systematic genetic and proteomic approaches with direct measurement of intracellular metals, we dissect nitroxoline perturbation of metal homeostasis and unveil additional effects on bacterial physiology. We show that nitroxoline affects outer membrane integrity, synergizing with large-scaffold antibiotics and resensitizing colistin-resistant Enterobacteriaceae *in vitro* and *in vivo*. We further characterise resistance mechanisms across *E. coli*, *Acinetobacter baumannii* and *Klebsiella pneumoniae*, recapitulating known *E. coli* resistance determinants and uncovering novel and conserved mechanisms across species, demonstrating their common effect on nitroxoline efflux.

## Introduction

Nitroxoline (8-hydroxy-5-nitroquinoline) is a quinoline-derivative (**Fig. 1a**) and FDA-approved antibiotic, used for more than 50 years as treatment and prophylaxis of acute and recurrent UTIs in several European and Asian countries^2–4^. Because of its excellent safety profile^3,5–7^ and activity against different organisms, nitroxoline has been recently proposed to be repurposed as antituberculosis^8^, antifungal^9,10^, antiviral^11,12^, antiparasitic^13^ and anticancer^14–17^ agent.

**Figure 1.**
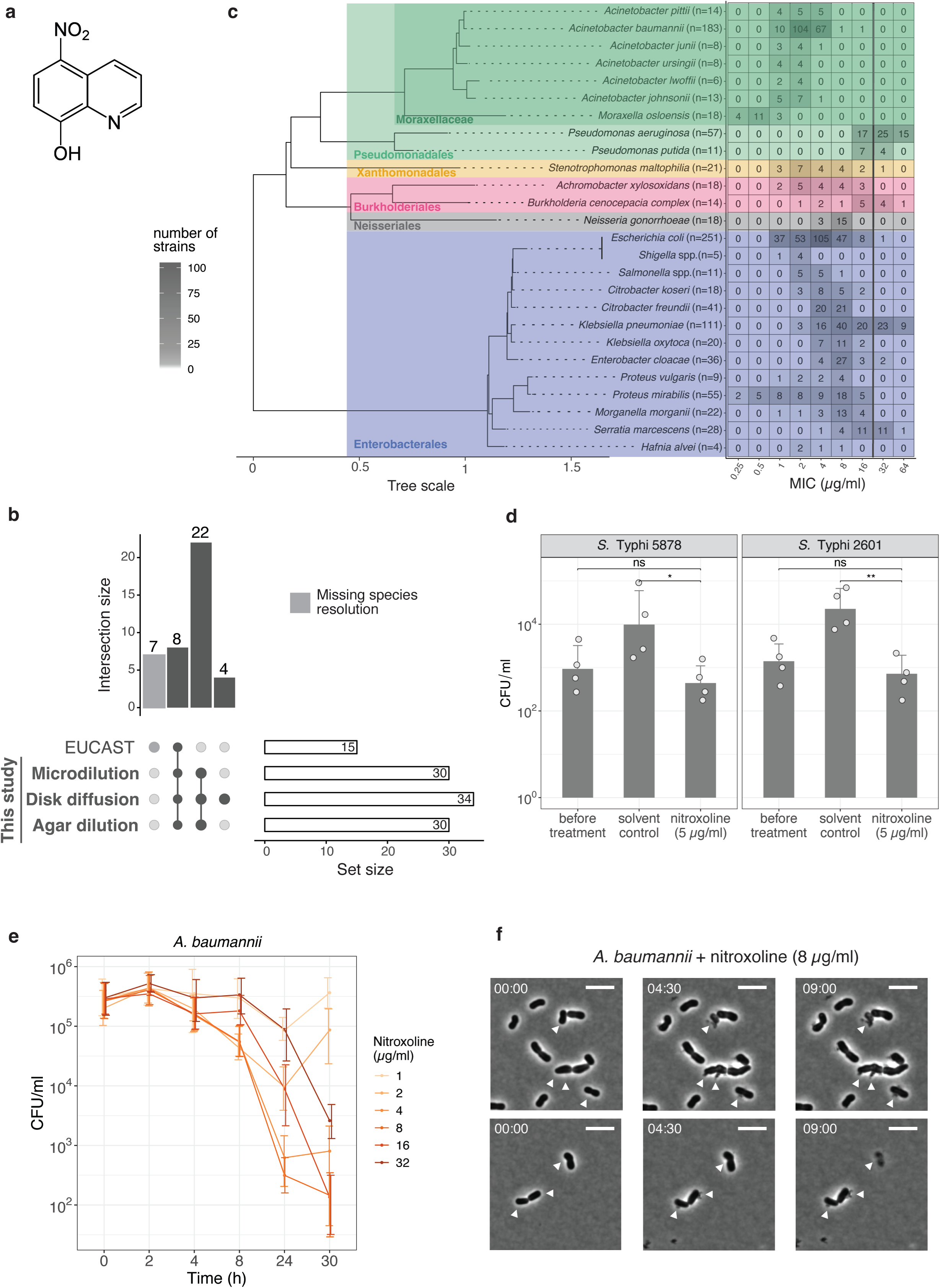
Nitroxoline is active beyond UTI pathogens, including intracellular bacteria, and exerts bactericidal activity. **a.** Nitroxoline structure. **b**. Overlap between Gram-negative bacterial genera or species tested with three orthogonal susceptibility testing methods in this study and according to EUCAST^19^. All seven EUCAST-specific entries are genera for which species resolution is missing. **c**. Nitroxoline is active against many Gram-negative bacterial species. MICs were determined against 30 bacterial species in broth microdilution (Methods). The total number of strains tested is indicated next to species, ordered by phylogeny according to GTDB^69^ (Methods), which classifies *E. coli* and *Shigella* as the same species. Multiple species are aggregated for *Salmonella* spp., *Shigella* spp., and *Burkholderia cenocepacia* complex. The clinical breakpoint for *E. coli* (16 µg/ml) is indicated (black line). **d**. Nitroxoline is active against intracellular *S.* Typhi. Intracellular bacterial counts were assessed with the gentamicin protection assay in two *S.* Typhi clinical isolates (Methods, **Supplementary Table 1**). Cell counts were determined before and after treatment with nitroxoline (5 µg/ml) or solvent control (DMSO) at 7 hours p.i. (MOI 100, 10 minutes incubation). Mean and standard error are shown across four independent experiments. ns p > 0.05; * ≤ p 0.05; ** p ≤ 0.01 (two-sided Welch’s *t*-test). **e.** Nitroxoline is bactericidal against *A. baumannii* ATCC 19606^T^ (Methods). Mean and standard deviation across at least three biological replicates are shown for each condition. The no-drug control is shown in Extended Data Fig. 2d. **f.** Nitroxoline induces lysis in *A. baumannii* ATCC 19606^T^. White arrowheads mark release of cytoplasmic material and loss of the pericellular halo. Representative images of phase-contrast videos were acquired after 8 µg/ml nitroxoline treatment (4 x MIC, Methods**, Supplementary Video 1**). Scale bar denotes 5 µm.

Nitroxoline has been shown to exert bacteriostatic activity against a broad range of Gram-negative and Gram-positive species^18,19^. Despite its long-standing use, information on nitroxoline PK/PD profile remains sparse^20^, with the only clinical breakpoint defined for *E. coli* from uncomplicated UTIs^2,21^. Metal chelation is assumed to underpin its general mode of action^22^, with reports on inhibition of biofilm formation^23,24^ and metallo-β-lactamases^25,26^. Nitroxoline has also been proposed to affect the activity of RNA-polymerase by chelation of Mg^2+^ and Mn^2+^ ^22^, and the activity of methionine aminopeptidases^5^. However, it remains unclear whether it acts as a metal chelator, sequestering essential metals, or as a metallophore, leading to metal stress, and which metals are differentially affected.

Nitroxoline resistance seems to be rare in *E. coli*^27^, with only a few resistance mechanisms identified to date *in vitro*. This includes the overexpression of the tripartite efflux pump EmrAB-TolC as a result of first-step mutations in the *emrR* gene (encoding a transcriptional repressor of the pump), and second-step mutations in *marR* and *lon*, conferring a higher *tolC* expression and resistance^28^.

Here we propose nitroxoline as a broader antibiotic against many Gram-negative species. We challenge previous knowledge of nitroxoline as a bacteriostatic agent, demonstrating bactericidal activity on pathogens for which therapeutic options are limited, such as *A. baumannii*. We uncover strong synergies of nitroxoline with several antibiotics, including colistin, resensitizing colistin-resistant Enterobacteriaceae *in vitro* and *in vivo*. Combining systems-based approaches with direct measurements of OM integrity and intracellular metal concentrations, we establish that nitroxoline acts as a zinc and copper metallophore and perturbs OM integrity. Finally, we show that the most recurring resistance mechanism across different pathogenic Enterobacteriaceae is the upregulation of Resistance-Nodulation-Division (RND) efflux pumps. Overall, we provide *in vitro* and *in vivo* evidence of nitroxoline’s ability to act, alone or in combination, against hard-to-treat bacterial pathogens, and provide further mechanistic understanding of its mode of action and resistance.

## Results

### Nitroxoline has a broad activity spectrum against Gram-negative bacteria

Although nitroxoline (**Fig. 1a**) has been used for decades against uncomplicated UTIs and has a good safety profile^3,5–7^, minimum inhibitory concentration (MIC) distributions are available from EUCAST only for eight bacterial species and seven genera^19^, with a clinical breakpoint determined only for *E. coli*^2^ (**Fig. 1b**). To investigate whether nitroxoline could be repurposed against other bacterial pathogens, we systematically profiled its susceptibility against 1,000-1,815 strains from 34 Gram-negative species. This set included some of the most relevant pathogens in the current antimicrobial resistance crisis, such as *A. baumannii*, *Burkholderia cenocepacia* complex and *Stenotrophomonas maltophilia* (**Fig. 1b-c**, **Extended Data Fig. 1a-b**).

We measured MICs in three orthogonal ways (**Fig. 1b**, **Methods**): broth microdilution (1,000 isolates, **Fig. 1c**), disk diffusion (1,815 isolates, **Extended Data Fig. 1a**) and agar dilution (1,004 isolates, **Extended Data Fig. 1b**). We observed good concordance between this study and EUCAST^19^ for the eight overlapping species (**Extended Data Fig. 1c**) and across the three methods, with the best agreement between broth and agar dilutions (Pearson correlation, R = 0.94) (**Extended Data Fig. 1d-f**).

From these results, we confirmed that several Gram-negative species were comparably or more susceptible to nitroxoline than *E. coli* (median MIC = 4 µg/ml for broth and agar dilution, lower than the EUCAST breakpoint of 16 µg/ml for *E. coli*^2^). For *E. coli* we confirmed this value, except for one strain out of 251 clinical isolates with MIC two-fold the breakpoint. Nitroxoline was active against Enterobacterales (median broth MIC: 4 µg/ml) and Moraxellaceae, which had the lowest MIC values (median broth MIC: 2 µg/ml) and comprise pathogens for which therapeutic options are currently limited, such as *A. baumannii*. Only *P. aeruginosa* exhibited a median MIC of 32 µg/ml above the clinical breakpoint (**Fig. 1c**). Overall, these results indicate nitroxoline as a promising antibacterial option against Enterobacterales and *Acinetobacter* spp.

### Nitroxoline is active against intracellular *Salmonella* Typhi and bactericidal against ***A. baumannii***

*Salmonella* spp. were among the most susceptible species to nitroxoline (average MIC in broth: 3.45 µg/ml; maximum MIC in broth: 8 µg/ml in only 1/11 tested strains) (**Fig. 1c**, **Extended Data Fig. 1a-b**). Since *Salmonella* can invade and persist in the host intracellularly^29^, we tested whether nitroxoline could also act on intracellular *Salmonella* Typhi, the leading serovar responsible for enteric fever^30^. Using an *in vitro* infection assay (gentamicin protection assay^31^, **Methods**), we showed that nitroxoline treatment results in a strong decrease (> 97% compared to solvent control) of intracellular *S.* Typhi in infected HeLa cells, for both clinical isolates tested (**Fig. 1d**).

As other metal chelators^32^, nitroxoline is classically considered bacteriostatic^33^. While we confirmed this in *E. coli* for concentrations eight-times its average MIC in broth (32 µg/ml) (**Extended Data Fig. 2a**), we detected partial bactericidal activity (decrease of at least 3 log_10_ colony-forming units (CFU) /ml at 24 hours^34^) against *A. baumannii*, for concentrations as low as 8 µg/ml, i.e. three times its average MIC of *A. baumannii* in broth (2.79 µg/ml) (**Fig. 1e**, **Extended Data Fig. 2b**, **Methods**). Cells released their cytoplasmic content and lysed as early as 4.5 hours of incubation with the drug (**Fig. 1f**, **Supplementary Videos 1-2**, **Methods**). To our knowledge, this is the first evidence of nitroxoline’s bactericidal activity.

### Nitroxoline antagonises beta-lactams and synergises with colistin

To explore potential combinatorial regimens, we tested nitroxoline in combination with 32 antibiotics in *E. coli* BW25113. The drug panel included all main classes of antibiotics used against Gram-negative bacteria, but also other metal chelators and antibiotics only effective against Gram-positive bacteria, such as vancomycin (**Methods**, **Supplementary Table 2**).

We uncovered extensive antagonism with beta-lactams (bactericidal cell-wall targeting drugs) including penicillins, cephalosporins and carbapenems (**Fig. 2a, Extended Data Fig. 3**, **Supplementary File**). Like other antagonisms between bactericidal and bacteriostatic drugs (such as nitroxoline at the concentration tested here), these interactions could be based on the fact that bactericidal drugs are more effective on actively dividing cells, and slowing down division with a bacteriostatic agent can alleviate their action^35^.

**Figure 2.**
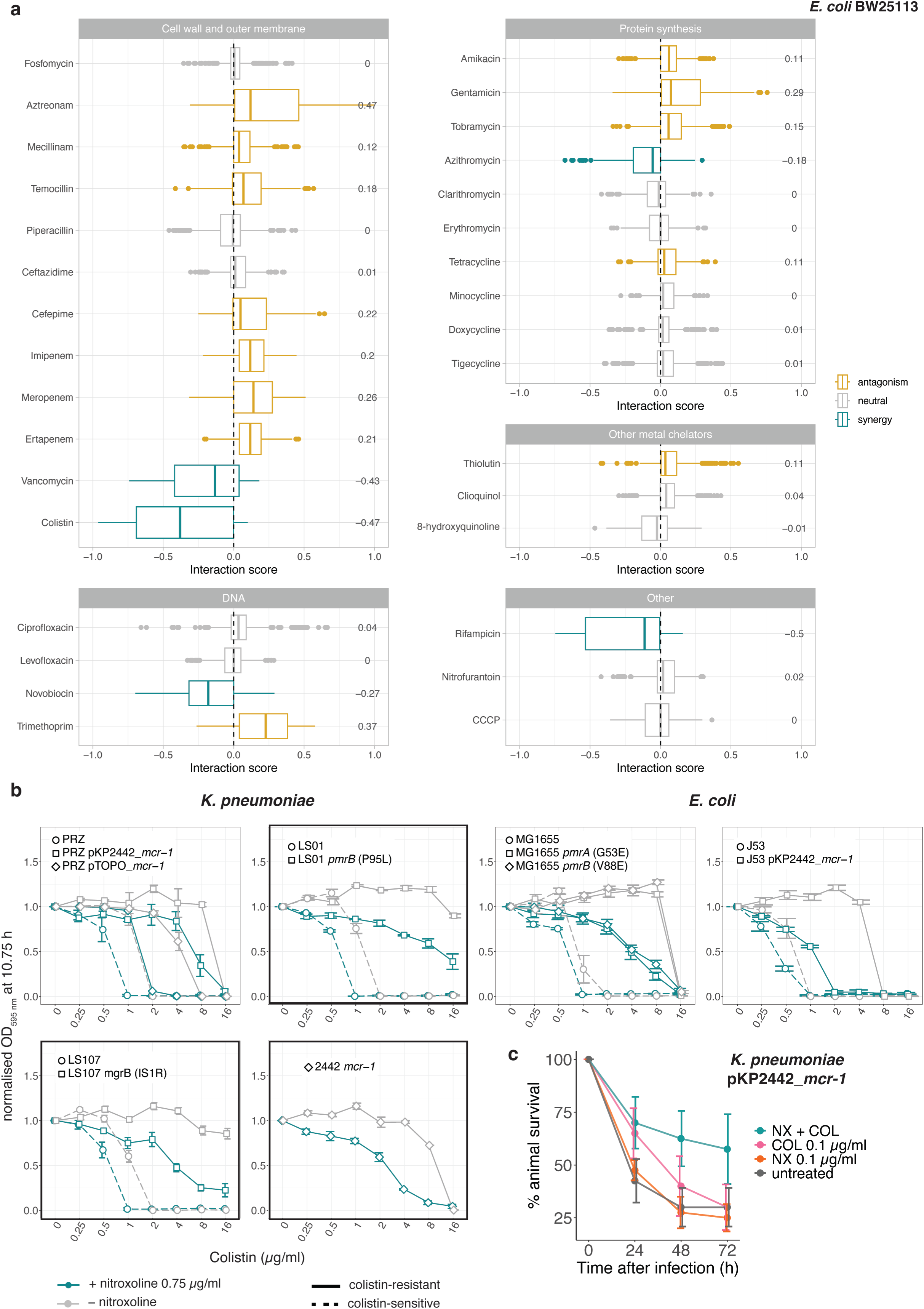
Nitroxoline interacts with other antibiotics in *E. coli* and resensitizes colistin-resistant *E. coli and K. pneumoniae*. **a.** Nitroxoline interacts with several antibiotics in *E. coli*. Nitroxoline combinations were tested in 8 x 8 broth microdilution checkerboards in *E. coli* BW25113 (**Extended Data** Fig. 3). Bliss interaction score distributions are shown for each combination. Median (central line), first (lower hinge) and third quartile (upper hinge) are shown for each boxplot. Whiskers correspond to 1.5 x IQR from each hinge. The numbers stand for cumulative Bliss scores for each combination (Methods). **b.** Nitroxoline resensitizes colistin-resistant *K. pneumoniae* and *E. coli* . Growth (endpoint OD_595nm_ corresponding to the beginning of stationary phase for the untreated control for each strain, Methods) was measured in the presence of serial two-fold dilutions of colistin, supplemented or not with 0.75 µg/ml nitroxoline and normalised by no-drug controls. Three *K. pneumoniae* and two *E. coli* strains (dashed lines) and their isogenic colistin-resistant descendants (solid lines) were tested, including experimentally evolved and clinical isolates (framed in black, Methods**, Supplementary Table 1**). One *K. pneumoniae* clinical isolate carries the *mcr-1* positive natural plasmid pKP2442 and therefore lacks a parental strain. Mean and standard error across four biological replicates are shown. Empty vector controls and full growth curves are shown in the Supplementary File. **c.** Nitroxoline resensitizes a colistin-resistant *K. pneumoniae* clinical isolate *in vivo*. *G. mellonella* larvae were infected with the indicated isolate and treated with single drugs or their combination. The percentage of surviving treated and untreated (solvent-only control) larvae was monitored over time. The mean and standard error are shown across four independent experiments for each condition. p = 0.0255 and p-0.0098 comparing colistin-nitroxoline with colistin and untreated, respectively (log-rank test). NX, nitroxoline; COL, colistin.

One of the most potent synergies of nitroxoline was with the OM-targeting drug colistin (**Fig. 2a**, **Extended Data Fig. 3**), whose toxicity limits its therapeutic use as last-resort agent^36^. Nitroxoline could therefore be used to lower colistin concentrations required to achieve therapeutic success, preventing toxicity. To explore this possibility, we tested whether nitroxoline could not only potentiate colistin action on sensitive strains, but also resensitize colistin-resistant strains. We showed that the addition of nitroxoline at sub-MIC concentration (0.75 µg/ml) decreases the MIC of *E. coli* and *K. pneumoniae* colistin-resistant strains (clinical and experimentally generated) from 2- to 4-fold, even below colistin EUCAST breakpoint (2 µg/ml)^2^ in three cases (**Fig. 2b, Supplementary File**). To confirm this synergy *in vivo*, we infected *Galleria mellonella* larvae with an *mcr-1* positive, colistin-resistant *K. pneumoniae* clinical isolate (**Methods**, **Supplementary Table 1**). The addition of nitroxoline improved the survival of the infected larvae by two-fold at 72 hours post-infection compared to the monotherapy with colistin or nitroxoline alone (**Fig. 2c**).

### Nitroxoline perturbs the OM in *E. coli*

In addition to colistin, nitroxoline synergised with all large-scaffold antibiotics, including drugs that are normally excluded by the OM and therefore are not active against Gram-negative bacteria, such as macrolides, rifampicin, novobiocin and vancomycin (**Fig. 2a, Extended Data Fig. 3**). Altogether, this suggested a direct effect of nitroxoline on the OM permeability.

To obtain a broader view on nitroxoline’s direct and indirect effects, we performed two-dimensional thermal proteome profiling (2D-TPP)^37^ on *E. coli* BW25113. Briefly, we exposed bacterial samples to multiple nitroxoline concentrations and subjected whole-cell or lysed samples to a gradient of temperatures, capturing nitroxoline effects on both abundance and stability of proteins (**Methods**). While changes in lysates will typically only detect direct target(s) of drugs, whole-cell changes provide a snapshot of both direct and indirect effects. We could not detect any significant change in lysate samples, suggesting that nitroxoline does not directly target a protein. In whole-cell samples we observed a decrease in the abundance/stability of OM proteins (OMPs) and members of LPS biosynthesis and trafficking (Lpt) machinery (**Fig. 3a**, **Extended Data Fig. 4a-b**, **Supplementary Table 3**).

**Figure 3.**
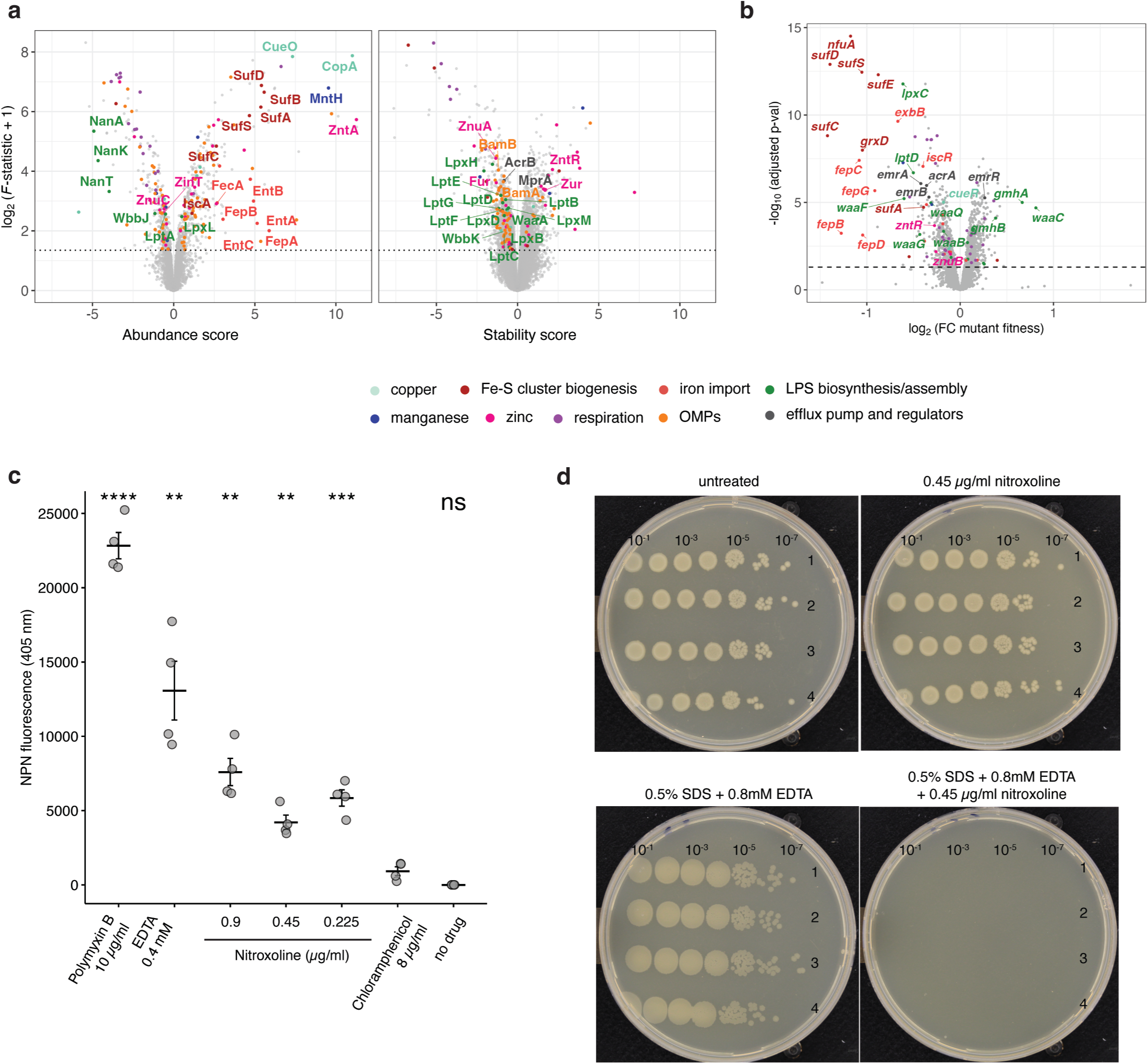
Nitroxoline directly perturbs the OM in *E. coli*. **a.** Nitroxoline decreases abundance and stability of outer membrane protein and Lpt machinery. Volcano plots depicting abundance or stability changes upon nitroxoline exposure in whole-cell 2D-TPP. Effect size and statistical significance (Methods) are represented on the x- and y- axis, respectively. For visualisation purposes, the F-statistic was linearly transformed to 1 when 0. Proteins are color-coded according to their annotation, derived from Gene Ontology (GO) (**Extended Data** Fig. 4a). **b**. Nitroxoline effects profiled by chemical genetics on an *E. coli* whole-genome single-gene deletion mutant library^39^. Effects are expressed as multiplicative changes of mutant fitness compared to the plate median (approximating wild-type) and significance was obtained from an empirical Bayes’ moderated t-statistics, Benjamini-Hochberg adjusted (Methods, **Supplementary Table 4**). Genes are color-coded as in Fig. 3a (GO in **Extended Data** Fig. 4b). **c.** Nitroxoline directly affects OM permeability. NPN fluorescence upon exposure of *E. coli* BW25113 to nitroxoline. Positive (polymyxin B, EDTA) and negative (chloramphenicol, untreated samples) controls are shown. Data points represent the average for each of four biological replicates per condition, across fluorescence measurements every 30 s over 10 minutes. The horizontal line and error bars indicate mean and standard error. ns p > 0.05; ** p ≤ 0.01; *** p ≤ 0.001; **** p ≤ 0.0001 (two-sided Welch’s *t*-test using the chloramphenicol control as reference group). **d.** Nitroxoline is more potent upon chemical perturbation of the OM. EOP assays with 10-fold serial dilutions of *E. coli* BW25113 cells plated onto no-drug control plates, 0.5% SDS–0.8 mM EDTA, 0.45 µg/ml nitroxoline, or a combination of the two conditions. Four biological replicates were tested for each condition. For all concentrations tested, including synergistic ones at which growth can be observed, see **Extended Data** Fig. 5c.

We combined this data with chemical genetics, in which we systematically mapped nitroxoline effects on the fitness of deletion mutants of every non-essential gene in *E. coli* ^38,39^. We found that mutants involved in similar processes, including LPS biosynthesis and transport, were more sensitive to nitroxoline, except for mutants involved in the first reactions of heptose incorporation into LPS inner core biosynthesis (*gmhA*, *gmhB* and *waaC*), which were more resistant (**Fig. 3b**, **Extended Data Fig. 4c, Supplementary Table 4**). This suggests the heptosyl-Kdo_2_ moiety of the lipopolysaccharide inner core as a minimum requirement for nitroxoline activity, since deletion of mutants catalysing most downstream LPS biosynthetic reactions, starting with *waaF*, are more sensitive to nitroxoline.

To provide direct evidence of nitroxoline’s effect on the OM, we quantified OM disruption using the hydrophobic probe 1-N-phenylnaphthylamine (NPN), which emits fluorescence upon exposure of the OM phospholipid layer^42^ (**Methods**). Sub-MIC concentrations of nitroxoline resulted in a significantly higher fluorescence than control samples (unexposed to any drug or to the non-OM-affecting antibiotic chloramphenicol). As positive controls, we used OM-targeting antibiotic polymyxin B and EDTA, another metal chelator that disrupts OM by sequestering LPS-stabilizing divalent cations^43^. Although lower than in positive controls, OM disruption by nitroxoline occurred even at 1/9 MIC (0.225 µg/ml, MIC = 2 µg/ml in *E. coli* BW25113, **Fig. 3c**).

To corroborate nitroxoline’s action on the OM, we tested its activity against the OM-defective *E. coli* strain *lptD4213*^40^ and in OM-perturbing conditions (0.5% SDS and 0.8 or 0.4 mM EDTA)^41^ using an efficiency of plating (EOP) assay (**Methods**). The *lptD4213* mutant was more susceptible to nitroxoline than wild-type *E. coli* (**Extended Data Fig. 4d**), in agreement with LptD decreased stability (**Fig. 3a**, **Extended Data Fig. 4b**) and with its loss-of-fitness already observed in the chemical genetic data, where the *lptD4213* mutant was included^38^ (**Fig. 3b**). Furthermore, nitroxoline synergized with the OM-perturbing conditions at concentrations 10-fold lower than MIC (**Fig. 3d**, **Extended Data Fig. 4e**).

It is possible that nitroxoline acts similarly as EDTA, chelating metals necessary for the stability of the OM^42^. While EDTA action is based on chelation of both Mg^2+^ and Ca^2+^, the two main cations involved in LPS stability^43^, nitroxoline has been shown to preferentially complex with Mn^2+^ and Mg^2+^, with no effect of Ca^2+^ supplementation on its MIC^22^. This could explain nitroxoline’s smaller effect than EDTA on OM integrity (**Fig. 3c**). However, the abundance of OMPs and Lpt machinery proteins was also altered (**Fig. 3a**, **Extended Data Fig. 4a-b**), suggesting an effect of nitroxoline on the regulation of the levels of these proteins.

### Nitroxoline acts as a zinc and copper metallophore

Nitroxoline is reported to chelate Mn^2+^ and Mg^2+^ and reach the intracellular milieu^22^, but a broader and more resolved view of its effects on metal homeostasis is missing. From the 2D-TPP (**Fig. 3a, Extended Data Fig. 4a**) and chemical genetic data (**Fig. 3b, Extended Data Fig. 4c**), we identified distinct profiles for proteins involved in the import, intracellular utilisation, and export of metals, consistent with responses to copper (**Fig. 4a**) and zinc (**Fig. 4b**) increase.

**Figure 4.**
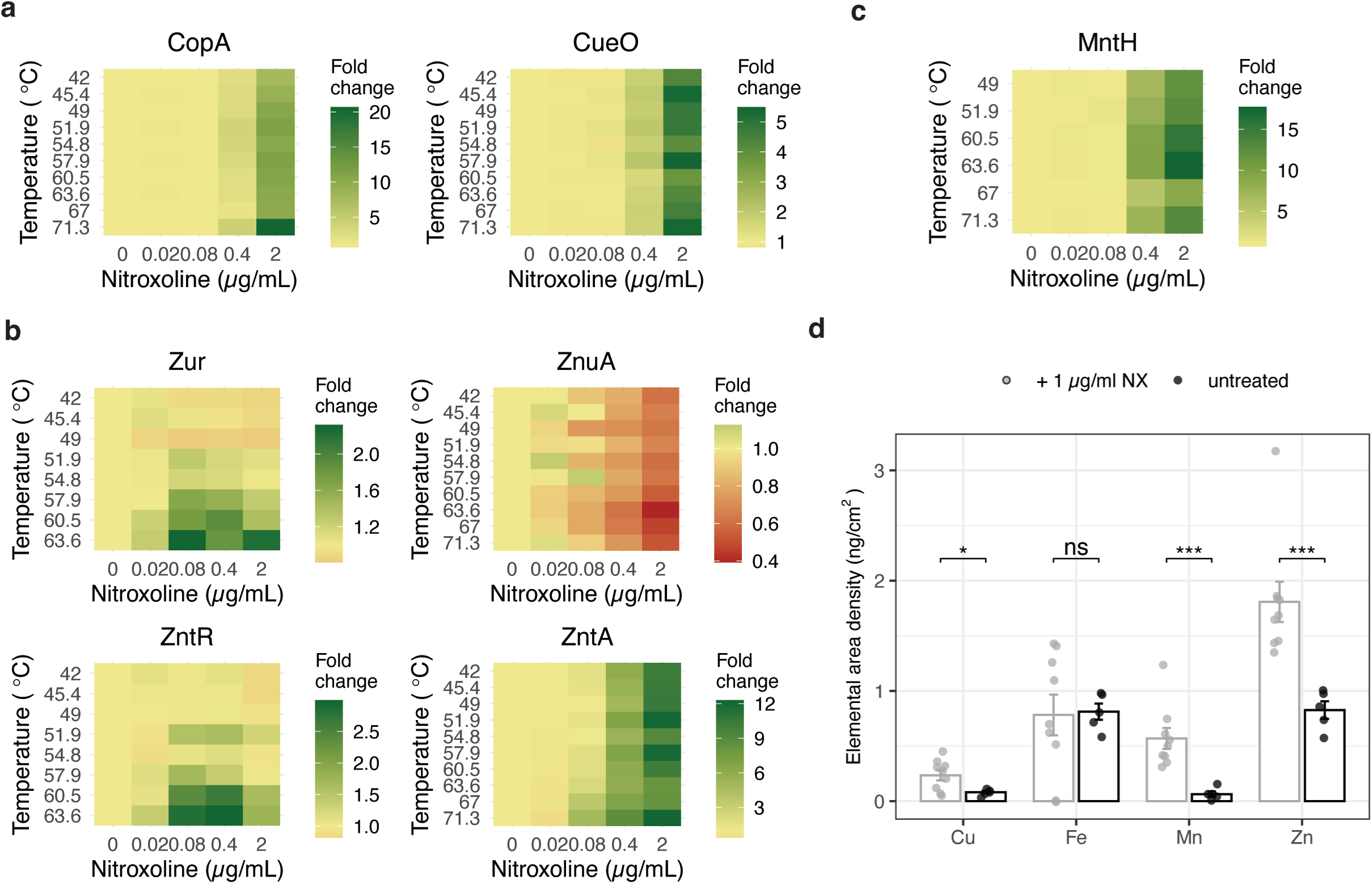
Nitroxoline increases intracellular levels of copper and zinc. **a-c.** Nitroxoline affects metal homeostasis inducing copper and zinc detoxification responses, as determined by 2D-TPP. Abundance and thermal stability profiles of the Cu(I) exporter CopA, the periplasmic copper oxidase CueO (**a**), the transcriptional regulators Zur and ZntR, zinc importer ZnuA and exporter ZntA (**b**), and the manganese importer MntH (**c**), are shown. **d.** Nitroxoline increases intracellular levels of copper, zinc and manganese. Synchrotron-based nano-XRF measurements on *E. coli* untreated or exposed to nitroxoline (1 µg/ml), expressed as elemental areal density (ng/cm^2^). The mean and standard error across ≥ 5 cells are shown (Methods). ns p > 0.05; * p ≤ 0.05; ** p ≤ 0.01 (two-sided Welch’s t-test).

At nitroxoline concentration around MIC (2 µg/ml), we observed an increased abundance of the copper-exporting P-type ATPase CopA (twenty-fold) and the multicopper oxidase CueO (five-fold), which upon Cu(I) excess remove copper from the cytoplasm and oxidise it in the periplasm, respectively^44^ (**Fig. 4a**, **Supplementary Table 3**). A similar effect has been reported for other quinolines, forming complexes with copper^45,46^, which may be extruded in the periplasm by CopA as a copper-overload defence mechanism. We observed similar effects for zinc, with the stabilisation of the zinc-responsive regulators Zur and ZntR and consistent changes in Zur- and ZntR-regulated proteins, such as subunits of the zinc importer ZnuABC (repressed by Zur) and the exporter ZntA (positively regulated by ZntR)^47^ (**Fig. 4b**). Accordingly, *ΔzntR* and *ΔznuB* were more sensitive to nitroxoline (**Fig. 3b, Supplementary Table 4**).

Zinc and copper intoxication has been associated with the disruption of iron-sulfur (Fe-S) clusters^48–51^ and a compensatory induction of iron-uptake proteins (derepressed by Fur), and Fe-S cluster biogenesis (*suf* genes, induced by IscR). Accordingly, upon nitroxoline exposure we observed decreased stability of Fur and an associated increase of Fur-repressed proteins involved in enterobactin biosynthesis (EntABF), recycling (Fes), receptor (FepA) and importing system (FepBCDG)^52^. The stability of IscR increased, with the associated decrease of *isc* operon and increase of *suf* operon members^53^ (**Fig. 3a, Extended Data Fig. 4f**). The increased abundance of the manganese importer MntH has also been reported as a consequence of copper stress^54^ (**Fig. 4c**). Since MntH is repressed by Fur, its increase is consistent with the observed Fur destabilisation (**Extended Data Fig. 4f**).

To confirm the impact of these effects on intracellular metal concentrations, we performed synchrotron-based nano-X-ray-fluorescence (XRF) on nitroxoline treated and untreated *E. coli*, confirming a four-fold copper, two-fold zinc, and ten-fold manganese increase in treated cells (**Fig. 4d**). Overall, our data suggest pleiotropic effects of nitroxoline on metal homeostasis, consistent with its activity as ionophore for copper, previously reported for clioquinol in cancer cells^46^, and zinc, as shown for other quinolines^55^.

### Nitroxoline resistance is based on conserved mechanisms across species

Our results suggest that nitroxoline does not have a direct protein target, but rather exerts pleiotropic effects on OM integrity and metal homeostasis, which might underpin the previously observed low frequency of resistance development^3,7,23,27,56^. To explore resistance mechanisms across different species, we evolved resistance to nitroxoline *in vitro* in three species: *E. coli*, for which nitroxoline is already used, and two Gram-negative species, *K. pneumoniae* and *A. baumannii*, for which nitroxoline could be repurposed considering its low MIC (**Fig. 1c**, **Extended Data Fig. 1a-b**).

We performed whole-genome sequencing (WGS) on 12 *E. coli*, 8 *K. pneumoniae* and 6 *A. baumannii* sensitive and evolved resistant strains (fold increase MIC ≥ 4 compared to parental sensitive strain) (**Methods**, **Fig. 5a, Supplementary Table 1**), and performed proteomics on a subset of them (**Fig. 5b**, **Extended Data Fig. 5a-b, Supplementary Table 5**). Mutations across species primarily affected transcriptional repressors of RND-type efflux pumps: *emrR* (previously reported in *E. coli*^28^), *oqxR* (*K. pneumoniae*), *adeL* and *tetR/acrR* (*A. baumannii*) (**Fig. 5a**).

**Figure 5.**
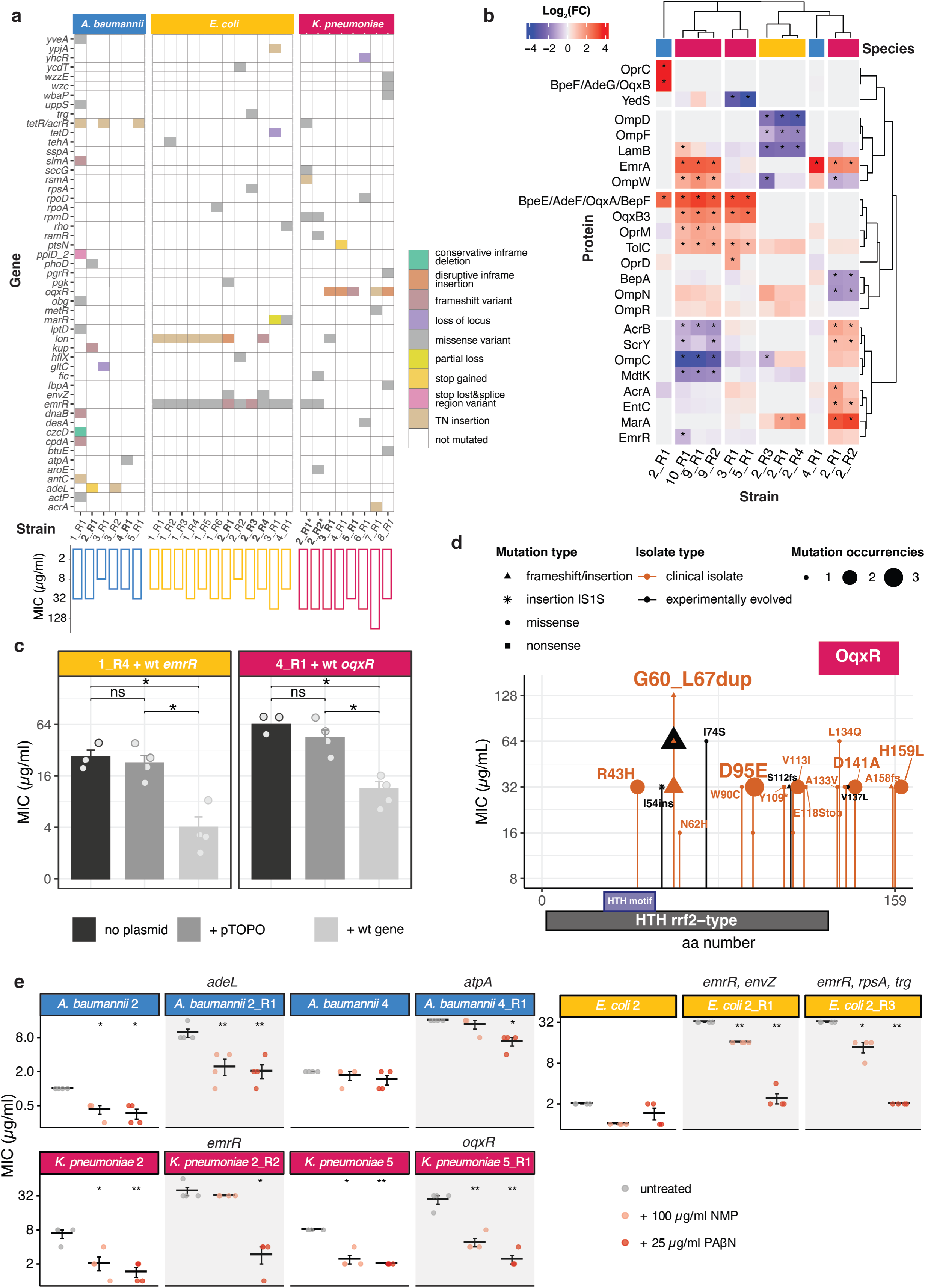
Resistance to nitroxoline is associated with efflux pump upregulation across species. **a.** Whole-genome sequencing of experimentally evolved nitroxoline-resistant strains (**Supplementary Table 1, 6**). Nitroxoline MIC values are indicated below each strain. Mutation effects are color-coded. Strains on which proteomics was performed (Fig. 5b) are indicated in bold. *K. pneumoniae* strains whose sensitive parental strain lacks *oqxR* are marked with an asterisk. The in-patient evolved *K. pneumoniae* clinical isolate 8_R1 is indicated in italics. **b.** Protein abundance changes in nitroxoline-resistant strains. Selected proteins annotated as efflux pumps or porins are shown and clustered according to Pearson’s correlation. Hits are marked with an asterisk (adjusted p-value ≤ 0.05 and at least two-fold abundance change, Methods). For all proteome changes of resistant strains see **Extended Data** Fig. 6a-b and **Supplementary Table 5**. Species are color-coded as in Fig. 5a. **c.** Wild-type *emrR* and *oqxr* complementation restores nitroxoline susceptibility. The experimentally evolved *E. coli* strain 1_R4 with *emrR* D109V mutation (Fig. 5a, **Extended Data** Fig. 6c) and *K. pneumoniae* strain 4_R1 harbouring the *oqxR* G60-L67 duplication (Fig. 5a, d) are shown. Nitroxoline MIC was measured by broth microdilution (Methods). **d.** Amino acid changes resulting from *oqxR* mutations. The domain annotation of OqxR was obtained from its closest annotated structural homolog NsrR (Methods). **e.** Efflux-pump inhibitors resensitize nitroxoline-resistant strains. Nitroxoline MIC was measured by broth microdilution (Methods). Resistant strains are shown as shaded plots next to their parental sensitive strains. For results on all strains see **Extended Data** Fig. 5g. p-values are only shown when significant: * p ≤ 0.05; ** p ≤ 0.01 (one-sided Welch’s t-test).

We found *emrR* mutations in all 12 evolved nitroxoline-resistant *E. coli* strains (**Fig. 5a**). This is consistent with previous reports^28^ and our chemical genetic data, where the *emrR* deletion mutant was more resistant and the knockouts of its regulated pump *emrAB* were more sensitive to nitroxoline (**Fig. 3b**). To verify the clinical relevance of these mutations, we assessed them in 14 clinical isolates with reduced susceptibility (MIC ≥ 8 µg/ml, i.e. at least two times the median MIC measured in this study for *E. coli*), finding distinct mutations from experimentally evolved strains (**Extended Data Fig. 5c**). We also found *emrR* mutations in two *K. pneumoniae* nitroxoline-resistant strains, which, as previously shown for *E. coli*^28^, had higher EmrA and TolC protein levels. An *A. baumannii* resistant strain, although lacking any mutation of efflux pump regulators, also exhibited a four-fold increase in EmrA levels (**Fig. 5b, Supplementary Table 5**).

Surprisingly, we did not detect any increase in EmrA and only a slight (< 2-fold) increase of TolC in three *E. coli emrR*-mutated strains, which showed instead a decreased abundance of porins OmpD, OmpF and LamB (**Fig. 5b**). For at least two of these strains (2_R1 and 2_R4), this could depend on missense mutations of *envZ*, that regulates porin expression via OmpR^57,58^, increased in these strains (**Fig. 5b**). Alternatively, porin abundance changes could be explained by mutations in the *lon* gene (**Fig. 5a**), previously associated with nitroxoline resistance^28^ and resulting in the stabilization of the Lon protease substrate MarA^59^. which regulates the expression of several drug resistance determinants, including porins^60,61^, and is also increased in these strains (**Fig. 5b**). Given the unexpected proteome changes in *emrR-* mutated *E. coli*, we sought to confirm the functional relevance of these mutations, complementing a strain carrying a recurring missense mutation (D109V, **Extended Data Fig. 5c**) with wild-type *emrR*, thereby restoring nitroxoline susceptibility (**Fig. 5c**).

The most frequent genetic alterations in *K. pneumoniae* resistant strains were mutations in *oqxR*, the transcriptional repressor of the OqxAB efflux pump, in agreement with recent reports^56^. We identified *oqxR* mutations in 5/8 experimentally evolved *K. pneumoniae* strains and in all 14 clinical isolates sequenced (**Fig. 5d**). We identified a mutational hotspot, common to clinical isolates and experimentally evolved strains: a duplication of eight amino acids (G60_67dup) resulting in a loop addition (**Fig. 5d, Extended Data Fig. 5d**). This mutation also emerged in a patient after a four-month prophylaxis with nitroxoline (*K. pneumoniae* urine isolate 8_R1, **Fig. 5a**), confirming its relevance for *in vivo* evolution of resistance. Accordingly, *oqxR*-mutated strains coclustered in the proteomics data (**Fig**. **5b****, Extended Data Fig. 5b**) and showed an increased abundance of OqxA (BepF), OqxB (OqxB3) and TolC (**Fig. 5b**, **Supplementary Table 5**). To further demonstrate the impact of this duplication on resistance, we complemented a G60_67dup-positive strain with wild-type *oqxR*, restoring nitroxoline susceptibility (**Fig. 5c**).

In resistant *A. baumannii* the most common mutations affected two transcriptional regulators: *adeL*, repressing the expression of the efflux pump AdeFG(BepF)-OprC, and a transcriptional regulator of the *acrR*/*tetR* family (**Fig. 5a**, **Extended Data Fig. 5e-f**). We performed proteomics on a strain carrying an *adeL* mutation resulting in a premature stop codon, which accordingly showed an increased abundance of all components of the efflux pump AdeFG-OprC (**Fig. 5b**). Another resistant strain, although not carrying any mutation in efflux pump regulators, exhibited a four-fold increase in EmrA abundance, which could explain its resistance (**Fig. 5b**).

From the mutational spectrum and proteomic changes that we observed across species, increased drug efflux via RND pumps appeared as a conserved strategy to achieve nitroxoline resistance. To verify this hypothesis, we tested the impact of the efflux pump inhibitors (EPI) 1-(1-naphthylmethyl)-piperazine (NMP) and phenylalanine-arginine β-naphthylamide (PAβN) on nitroxoline susceptibility. We observed a decrease in nitroxoline MIC both in resistant and susceptible strains, independent of their specific mutations and generally more marked for PAβN, previously reported as an inhibitor of RND pumps in *E. coli*^62^ and of AdeFG in *A. baumanni*^63^ (**Fig. 5e**, **Extended Data Fig. 5g**, **Methods**).

## Discussion

The alarming spread of antimicrobial resistance is aggravated by the slow development of new compounds. This is not only due to the experimental challenge of developing novel compounds, ideally with novel bacterial targets and low resistance potential, but also to economic hurdles in bringing new compounds to the clinic. In this context, repurposing already approved drugs, with known PK/PD and toxicity profile, holds great potential to accelerate the clinical translation of novel antibacterial strategies.

With nitroxoline, we show how revisiting the spectrum and mode of action of an FDA-approved drug opens new therapeutic possibilities for some of the most challenging bacterial species in the current AMR scenario like colistin-resistant Enterobacteriaceae and *A. baumannii*. In addition to its usage as single drug, we demonstrate nitroxoline as a powerful synergizer in combination with other drugs, sensitizing *E. coli* to antibiotics normally bottlenecked by the OM and active only against Gram-positive species. Additionally, nitroxoline resensitized colistin-resistant Enterobacteriaceae, independently of the species and the colistin resistance determinant, including *in vivo* against an *mcr-1* positive *K. pneumoniae* clinical isolate. While further studies are needed to verify nitroxoline’s adequate therapeutic concentrations beyond its current UTI indications, our results, as well as recent anti-cancer formulations for other anatomical regions than the bladder^15,64,65^, suggest that the activity that we demonstrate *in vitro* and *in vivo* could also be achieved in humans for novel therapeutic uses.

Despite its decade-long use, the mode of action of nitroxoline has remained elusive. Here, we revisit its activity with systems-biology approaches, such as chemical genetics, 2D-TPP and high-throughput drug combinatorial testing. The integration of this data uncovered new effects, such as nitroxoline’s action on the OM, on the abundance of OMPs and members of the Lpt machinery (**Fig. 3a-b, Extended Data Fig. 4a-e**).

Nitroxoline has classically been considered a metal chelator: using the 2D-TPP and chemical genetics and directly measuring intracellular metals, we propose a novel activity as metallophore, causing copper and zinc accumulation in the cell, and thereby metal stress/toxicity. This hypothesis is futher supported by the conserved physiological responses across various species upon nitroxoline exposure, such as the increase of the copper and zinc exporters, of siderophores (likely as a response to the damage of FeS clusters by metal stress), and of phenols and polyamines, known to act as antioxidants particularly upon metal intoxication (**Fig. 3a-b, 4a-c, Supplementary Fig. 7-8, 9a**).

Considering our results, the broad antibacterial spectrum against Gram-negative bacteria could be attributed to the fact that nitroxoline (i) seems to lack a specific protein target, (ii) has pleiotropic effects on the OM and OMP levels and (iii) acts as an ionophore inducing intracellular metal intoxication for zinc and copper. We highlighted critical species-specific differences in nitroxoline’s mode of action, such as its bactericidal activity in *A. baumannii*, challenging the definition of nitroxoline as a bacteriostatic agent, and pointing towards envelope damage. Since drugs can be bacteriostatic or bactericidal depending on the strain considered^66^, future studies should explore the conservation of such activity across multiple *A. baumannii* strains and its mechanistic underpinnings.

We also revealed cross-species mechanisms for resistance, such as the regulation of efflux pumps, part of the RND superfamily prevalent in Gram-negative bacteria. A few shared responses were previously observed for another quinoline, chloroxine^45^, including the increase of MarA, also resulting in efflux pump upregulation or porin downregulation (as we showed for *E. coli*) and of the nitroreductases NfsA and NfsB. While this could suggest cross-resistance between nitroxoline and nitrofuran antibiotics, for which this is the most common resistance determinant, this has been previously disproven at least in *E. coli*^28^. Importantly, neither through resistance evolution nor in naturally occurring resistant isolates could we detect mutations on potential protein targets, in concordance with previous reports^28^ and with our 2D-TPP results, which did not identify any protein stabilization in lysates. This supports the absence of a specific protein target for nitroxoline and excludes, to the best of our knowledge, an important potential resistance mode.

In summary, we show how revisiting a compound used for decades with systems approaches can reveal a novel spectrum, mode of action and resistance mechanisms, offering new and safe therapeutic possibilities against hard-to-treat bacterial species.

## Methods

### Bacterial strains and growth conditions

All strains used in this study are listed in Supplementary Table 1. Unless otherwise specified, bacteria were grown in cation-adjusted Müller-Hinton (MH II) broth at 37 °C with continuous shaking at 180 rpm in 5 ml for overnight cultures and in 50 µL in microtiter plates. For growth on solid medium, MH II was supplemented with 1.5% agar.

### MIC measurement and efflux pump inhibitor supplementation

Antimicrobial susceptibility was determined by disc diffusion (Kirby Bauer assay), agar dilution (nitroxoline concentration range: 0.125-128 µg/ml) and broth microdilution (range: 0.125-64 µg/ml) as previously described^67^. MICs and zone of inhibitions were evaluated and interpreted according to EUCAST^2^. Efflux pumps, 1-(1-Naphthylmethyl)-piperazine (NMP) and Phenylalanine-Arginine β-naphthylamide dihydrochloride (PAβN) were diluted in 25 g/L stock solutions in DMSO and used as previously described^68^.

### Species phylogeny analysis

A phylogeny tree was constructed from the Genome Taxonomy Database (GTDB) bacterial reference tree (release 08-RS214^69^) using the ETE toolkit^70^. The GTDB taxonomy decorating the tree was then used to convert genome IDs to their corresponding species names. The tree was visualized using the R package ggtree^71^.

### Time-kill curves

A bacterial suspension in 0.9% NaCl (McFarland standard of 0.5) was prepared from overnight cultures, diluted 1:100 in 10 ml of MH II broth and incubated at 37 °C with continuous shaking for 30 h with a two-fold dilution series of nitroxoline (1/2-16x MIC), or DMSO as no-drug control. 100 µL of cells were collected at specified time intervals, serially diluted in PBS (10^0^ to 10^-9^ dilutions) and spread onto blood agar plates. Cell viability was determined by counting colony-forming units (CFUs).

### *A. baumannii* time-lapse imaging

Cells were grown overnight, diluted to an OD_600nm_ of 0.01 and grown for 3 h at 37 °C, as described in the “Growth conditions” section. Cells were then spotted on MHII + 1% agarose pads, supplemented or not with 8 μg/ml of nitroxoline (four-fold MIC) between a glass slide and a coverslip. Slides were sealed with Valap to avoid coverslip shifting. Imaging was performed at room temperature every 15 minutes for 10 hours over three distinct points of the slides for each condition using a Nikon Eclipse Ti inverted microscope, equipped with a Nikon DS-Qi2 camera, a Nikon Plan Apo Lambda ×60 oil Ph3 DM phase-contrast objective. Images were acquired with NIS-Elements AR4.50.00 software and processed with Fiji v.2.9.0/1.53t.

### Gentamicin protection assay and quantification of intracellular *Salmonella*

HeLa cells were cultivated in cell culture flasks (75 cm^2^) with RPMI medium. One day prior to infection, cells were seeded into 24-well plates (10^5^ cells per well). *S.* Typhi clinical isolates (**Supplementary Table 1**) were grown overnight in LB medium and subcultured (1:33) for three hours at 37 °C with shaking. Bacteria were harvested (13,000 x *g*, 5 min), resuspended in RPMI medium and subsequently used for infection at a multiplicity of infection (MOI) 100 for 10 minutes. After discarding the supernatant, cells were washed with PBS, incubated for 40 minutes in RPMI with 100 µg/ml gentamicin to eliminate extracellular bacteria. Cells were then washed twice with PBS and incubated with RPMI with 5 µg/ml nitroxoline (previously tested to exclude toxicity on HeLa cells alone) or DMSO as a vehicle control for 7 hours. Infected cells were washed with PBS, lysed with 1% Triton-X and 0.1% SDS, and serial dilutions in PBS (10^0^ – 10^-3^) were plated on LB agar. Plates were incubated overnight at 37 °C and bacterial colonies were counted in at least four independent experiments in two technical replicates.

### Concentration pretesting for checkerboard assay and data analysis

12 two-fold serial dilutions of nitroxoline and 32 other compounds (**Supplementary Table 2**) were arrayed in technical duplicates in 384-well plates (ref. 781271 by Greiner BioOne) and inoculated with *E. coli* K-12 at a starting OD_600nm_ of 0.001. Plates were sealed with breathable membranes incubated at 37 °C with continuous shaking and OD_600nm_ was measured every 30 minutes for 14 hours. The background, corresponding to the OD_600nm_ at the first time point, was subtracted from each measurement for each well. The point at the transition from exponential to stationary phase was detected for each well and corresponding OD_600nm_ normalised by the median of the corresponding value of the no-drug controls present in each plate (n = 16). For each drug, the IC75, i.e. the concentration at which 75% of the growth was inhibited, was identified in the resulting dose-response curves for each drug. For the checkerboard assay, eight evenly spaced concentrations were then selected, with the highest one corresponding to the IC75 and the lowest one corresponding to the no-drug control. Nitroxoline was tested in almost all combinations at IC50 to be ideally placed to discover synergies as potential combinatorial regimens. All experiments were conducted in biological duplicates (i.e. plates inoculated with overnight cultures from distinct colonies).

### Checkerboard assay and data analysis

Pairwise combinations of nitroxoline with 32 compounds (**Supplementary Table 2**) were tested in a checkerboard microdilution assay. Drugs were arrayed in 8 x 8 checkerboards using the eight concentrations previously selected. Growth was measured in the same conditions and data was analysed as for the concentration pretesting. The OD_600nm_ at the transition between the exponential and stationary phase, after background subtraction, was normalised by the median of the corresponding value of the no-drug controls present in each plate (n = 6). This value was used to calculate Bliss interaction scores ε ^72^ for each drug-drug concentration combination as follows:

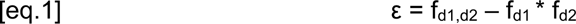

where f_d1d2_ corresponds to the observed fitness in the presence of the drug combination, and f_d1_ and f_d2_ correspond to the fitness in the presence of the two single drugs. We therefore obtained 49 ε scores for each checkerboard replicate. All experiments were conducted in at least two biological replicates, resulting in at least 98 ε scores for each combination. Synergies and antagonisms were assigned when the first and third quartile of the ε distribution, respectively, exceeded |0.1| and the median ε exceeded |0.03|. Cumulative Bliss scores for each combination were considered as first quartile, third quartile and median, for synergies, antagonisms and neutral interactions, respectively.

### Resensitization of colistin-resistant strains by nitroxoline

Cells were pre-cultured as in “Growth conditions”, growth was measured and data was analysed as in “Concentration pretesting for checkerboard assay and data analysis” in plates containing eight two-fold dilutions of colistin, supplemented or not with nitroxoline at 0.75 µg/ml. The strains used are listed in Supplementary Table 1.

### Two-dimensional thermal proteome profiling (2D-TPP)

Cells were grown overnight, diluted 1000-fold and grown until OD_578nm_ ∼0.6 at 37 °C, as described in the “Growth conditions” section. After addition of nitroxoline at the selected concentrations (0.02, 0.08, 0.4 and 2 µg/ml) or a vehicle-treated control, cultures were incubated at 37 °C for 10 minutes. After 4,000 x *g* centrifugation for 5 min, cells were washed with 10 ml PBS containing the drug at the appropriate concentrations and resuspended in the same buffer to an OD_578nm_ of 10. 100 µL of this suspension was then aliquoted to ten wells of a PCR plate that was centrifuged at 4,000 x *g* for 5 min. 80 µL of the supernatant was removed before exposing the plate to a temperature gradient for 3 minutes in a PCR machine (Agilent SureCycler 8800), followed by 3 minutes at room temperature. Cells were lysed with 30 µL lysis buffer (final concentration: 50 μg/ml lysozyme, 0.8% NP-40, 1x protease inhibitor (Roche), 250 U/ml benzonase and 1 mM MgCl_2_ in PBS) for 20 minutes, shaking at room temperature, followed by three freeze–thaw cycles. Protein aggregates were removed by centrifuging the plate at 2,000 x *g* and filtering the supernatant at 500 x *g* through a 0.45 µm filter plate (Millipore, ref: MSHVN4550) for 5 minutes at 4 °C. Protein digestion, peptide labelling, and MS-based proteomics were performed as previously described^73^.

### 2D-TPP data analysis

Data were pre-processed and normalised as previously described^74^, using the *E. coli* K-12 strain Uniprot FASTA (Proteome ID: UP000000625), modified to include known contaminants and the reversed protein sequences, to perform peptide and protein identification. Data analysis was performed using the R package TPP2D^75^. In brief, a null-model, assuming that the soluble protein fraction depends only on temperature, and an alternative model, assuming a sigmoidal dose-response function for each temperature tested, were fitted to the data. For each protein, an F-statistic was obtained from the comparison of the residual sum of squares (RSS) of the two models. Abundance or thermal stability effect size were calculated for each protein as:

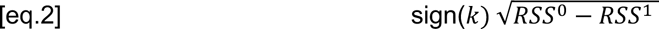

where k is the slope of the dose-response model fitted across temperatures and drug concentrations, RSS^0^ and RSS^1^ correspond to the residual sum of squares of the null (pEC50 linearly scaling with temperature) and alternative model, respectively^75^.

### Gene Ontology (GO) enrichment

The enrichment analysis was performed on proteomes of *E. coli* BW251113 for the 2D-TPP data and for the proteomics data, and of the strains listed in Supplementary Table 1 as used for “nitroxoline resistance evolution” for the analysis of proteomics data on sensitive and resistant strains. Proteomes were annotated using GOs downloaded from http://geneontology.org/ (release 2022-11-03). For each GO term, the enrichment of input protein sets (hits corresponding to FDR < 0.05) against the background (all detected proteins) was tested using Fisher’s exact test. P-values were corrected for multiple testing using the Benjamini-Hochberg procedure.

### Nitroxoline MIC in *E. coli lptD4213*

Cells were grown as in “Growth conditions”. Growth was measured as in “Concentration pretesting for checkerboard assay and data analysis” upon exposure to seven nitroxoline two-fold dilutions in *E. coli* BW25113 and *E. coli lptD4213*. Data was analysed as in “Resensitization of colistin-resistant strains by nitroxoline”. Experiments were conducted in three biological replicates.

### Evaluation of drug combination therapy using the *G. mellonella* infection model

Larvae of the greater wax moth (*Galleria mellonella*) were infected with *Klebsiella pneumoniae* and treated with single drugs or drug combinations as previously described^76^. Caterpillars were purchased from Valomolia (Strasbourg, France). Stock solutions of colistin and nitroxoline were freshly prepared with 20 mM sodium acetate buffer (pH 5). Drug toxicity was preliminarily determined by injecting larvae with serial dilutions of single drugs and combinations. Non-toxic concentrations of the drugs were used for further experiments. Bacteria were grown overnight as described in **“**Growth conditions**”**, harvested at an OD_600nm_ of 0.2, washed with PBS and adjusted to 5.6×10^7^ cfu/ml corresponding to a median lethal dose of 60-70% after 24 h, as determined in preliminary experiments. Groups of 10 caterpillars per condition were injected with 10 µL of the bacterial suspension into the haemocoel via the last right proleg and incubated at 37 °C. One hour post-infection caterpillars were injected into the last left proleg with 10 µL of single drugs or drug combinations (0.1 µg/ml colistin and 0.1 µg/ml nitroxoline). Survival was monitored for 72 h. Each strain–drug combination was evaluated in three independent experiments. The statistical analysis was performed using the log-rank test.

### Efficiency of plating (EOP) assay

Cells were grown overday for 8 hours as in “Growth conditions” and ten-fold serially diluted eight times. From each dilution, 3 µL were spotted onto MH II plates supplemented or not with 0.8 mM EDTA–0.5% SDS, 0.45 µg/ml nitroxoline, and a combination of the two conditions. Spots were allowed to dry and the plates were incubated overnight at 37 °C. Experiments were conducted in four biological replicates for each condition.

### *N*-phenylnaphthylamine (NPN)-fluorescence assay for OM damage

The assay was conducted as previously described^77^. Briefly, cells were grown overnight as described in the “Growth conditions” section and diluted to an OD_600nm_ = 0.5 in 5 mM pH 7.2 HEPES buffer (Sigma Aldrich). 100 µl of the cell suspension, together with 50 µl drugs diluted in HEPES buffer at the appropriate concentrations and 50 µl *N*-phenyl-1-naphthylamine (NPN) diluted in HEPES to a final concentration 10 µM, were added to a black 96-well plate with clear-bottomed wells. Controls included all possible combinations of cells, drugs and NPN, each of them separately, and a plain buffer control. Fluorescence was measured immediately on a Tecan Safire2 plate reader using an excitation wavelength of 355 nm and an emission wavelength of 405 nm. Fluorescence measurements were obtained every 30 s for 10 min. After averaging across the 20 replicated measurements, the NPN Uptake Factor was calculated as follows:

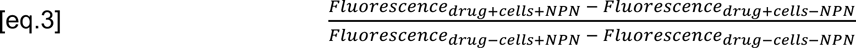

Finally, the uptake values of samples containing drugs were compared to the no-drug control. Positive controls included 10 µg/ml polymyxin B and 0.4 mM EDTA. As a negative control, a non-OM-perturbing antibiotic (chloramphenicol) at its MIC (8 µg/ml) was included. Because of quenching between the NPN emission wavelength and nitroxoline excitation wavelength, nitroxoline concentrations higher than 2 µg/ml showed a linear decrease in fluorescence and were not used. Experiments were conducted in four biological replicates.

### Measurement of metal abundance and distribution via synchrotron radiation-induced X-ray fluorescence nano-imaging

Experiments were performed at the Nano-imaging beamline ID16A of the European Synchrotron Radiation Facility (ESRF). *E. coli* BW25113 cells were grown overnight in LB as described in “Growth conditions”, subcultured until reaching OD_600nm_ 0.1 and treated for 15 minutes with nitroxoline (1 µg/ml) or 0.1% DMSO at 37 °C. After washing twice in PBS, 10 µL were mounted on silicon nitride membranes (Silson, Southam, UK, 1.5 mm × 1.5 mm × 0.5 µm) before cryo-fixation using a freeze plunger (EM GP, Leica) with 1 s blotting time. A 17 keV X-ray beam was focused to a 45 nm horizontal × 37 nm vertical spot by a pair of multilayer-coated Kirkpatrick-Baez mirrors located 185 meters downstream of the undulator source with a high flux of 4.1×10^11^ ph/s. The samples were rastered through the focal spot of the beam under a vacuum of 10^-7^ mbar at −179 °C. XRF spectra were measured using a pair of element silicon drift diode detectors (7-element detector Vortex-ME7, Hitachi, and 16-element detector) to subsequently quantify elements by their K-level emission lines. Low-resolution and fast-position mapping by combined X-ray phase contrast and XRF coarse scans in low-dose mode (6.1×10^10^ ph/s) were performed using a scan step size of 300 × 300 nm^2^ and a dwell time of 100 ms to identify bacteria. XRF fine scans in high-dose mode (2.49×10^11^ ph/s) were then performed with a step size of either 30 × 30 or 15 × 15 nm^2^ and a dwell time of 50 ms to obtain quantitative elemental density maps from individual point spectra. After fitting and normalizing the data with PyMca XRF spectral analysis software, mean intracellular elemental area density (ng/mm^2^) were calculated from at least two different areas from two independent experiments.

### *E. coli* chemical genetic screen

Nitroxoline was tested on the *E. coli* whole-genome single-gene deletion mutant Keio collection^39^ as previously described^38^. The collection (two independent clones per mutant), which was cryopreserved in 384-well plates, was arrayed in 1536-colony format using a Rotor HDA (Singer Instruments). Cells were grown at 37 °C for 10 hours and pinned on LB-plates, with or without nitroxoline (2 µg/ml) in three replicates. After 16 hours of incubation at 37 °C, plates were imaged using a controlled-light setup (spImager, S&P Robotics) and an 18-megapixel Canon EOS Rebel T3i camera. Mutant growth was calculated by quantifying colony opacity, estimated with the Iris software^78^. To account for the better growth at the edges of a plate, two outermost columns/rows were multiplicatively adjusted to the median opacity of the plate^79^. Mutant fitness was then estimated as a fraction of the plate median opacity. A change in mutant fitness was quantified as a multiplicative change per condition using a two-sided unpaired t-test. The resulting t-statistic was empirical Bayes’ moderated^80^ and corresponding p-values were adjusted for multiple testing (Benjamini-Hochberg correction^81^) (**Supplementary Table 4**).

### Experimental resistance evolution

Nitroxoline-sensitive clinical isolates and reference strains of *E. coli*, *K. pneumoniae* and *A. baumannii* (**Supplementary Table 1**) were exposed to increasing nitroxoline concentrations from 0.5x MIC to 4x MIC (two-fold dilution steps) in LB as previously described^28^. A defined bacterial inoculum (McFarland 0.5, corresponding to 10^8^ bacteria/ml) was passaged every 24 hours for at least 7 days in 5 ml LB. In case of bacterial growth, nitroxoline concentration was increased two-fold. The MIC of the strains was measured using broth microdilution and agar dilution as described before.

### Genome sequencing and single nucleotide polymorphism (SNP) analysis

Experimentally evolved strains were defined as resistant if MIC fold increase ≥ 4 compared to parental strain. Clinical isolates were considered resistant if their MIC was at least two times the median species MIC measured in this study (**Fig. 1c, Extended Data Fig. 1a-b**). Genome sequencing was performed for all isolates using short-read technology (MiSeq platform; Illumina, San Diego, CA) generating 150 or 250 bp paired-end reads and >100-fold average coverage. After quality trimming of the reads, *de novo* assembly and scaffolding was conducted using SPAdes version 3.12.0 with standard parameters. Annotation was done with Prokka version 1.14.6^82^. SNP analysis was performed using snippy (https://github.com/tseemann/snippy) to compare isogenic nitroxoline susceptible and resistant strains. Deletions were analysed using an in-house script. Clinical isolates were further compared to annotated reference genomes (GCF_000258865.1, GCF_000750555.1 and Bioproject PRJNA901493) to determine mutations in genes encoding for efflux pump regulators with an in-house script.

### Structural alignment and annotation of OqxR, EmrR, AdeL, AcrR/TetR

Amino acid changes in mutated proteins from nitroxoline-resistant clinical isolates were mapped onto protein features extracted from Proteins API^83^ using a custom-script, adapted from the software mutplot^84^ (UniProt IDs used: EmrR: P0ACR9; AdeL: A0A059ZJX1; AcrR/TetR: A0A245ZZS0). Because OqxR in *K. pneumoniae* lacks a UniProt ID and feature annotation, we searched for its closest, feature-annotated, structural homolog with Foldseek using the 3Di/AA mode^85^. The highest-ranking protein (E-value 7.58E-07, score 265) with available domain annotation was another Rrf2 transcription factor, NsrR from *S. enterica subsp. enterica* serovar Typhimurium LT2 (AF-Q8ZKA3-F1-model_v4). Structures were visualized using Mol* Viewer^86^ from PDB^87^. Alignment of wild-type and mutated OqxR structures were performed using the Pairwise Structure Alignment tool on RCSB PDB^88^. Amino acid sequences of the wild-type and mutated forms can be found in the **Supplementary File**.

### Complementation of nitroxoline resistant isolates

Clinical isolates with *emrR* or *oqxR* mutations were transformed with pTOPO expression plasmids (pCR-Blunt II-TOPO, Invitrogen) harboring the wild-type gene (pTOPO_*emrR* or pTOPO_*oqxR*) via electroporation as previously described^89^. For this purpose, bacteria were grown over night as described in “Growth conditions”. On the next day, cells were subcultivated (1:100 dilution) until an OD_600nm_ of 0.4-0.6 was reached. Cells were harvested (13,000 x *g*, 3 min), washed once in 500 µL 300 mM ice-cold sucrose, and transformed with 500 ng of plasmid DNA with a Gene Pulser Xcell electroporator (Bio-Rad) with 2.2 kV, 200 Ω, and 25 µF settings. Cells were recovered in SOC medium at 37 °C for one hour (with shaking at 180 rpm) and plated on LB agar plates with kanamycin (30 mg/L for *E. coli* and 100 mg/L for *K. pneumoniae* and *A. baumannii*) for selection of transformants, subsequently used for MIC testing.

### Proteome profiling of mutant strains

Cells were grown overnight as described in “Growth conditions”, diluted to OD_600nm_ = 0.05 in 3 ml LB, grown until reaching OD_600nm_ = 0.5. Nitroxoline-sensitive strains were treated with 1x MIC nitroxoline (0.5-4 µg/ml depending on the strain) or with the DMSO control for 10 min. Nitroxoline-resistant strains were exposed only to DMSO. After 4,000 x *g* centrifugation for 5 min, 2 ml aliquots were washed with 1 ml PBS (containing the drug at the appropriate concentration for drug-exposed samples). The final pellets were frozen at −20 °C until analysis, when they were resuspended in lysis buffer (final concentration: 2% SDS, 250 U/ml benzonase and 1 mM MgCl_2_ in PBS) and immediately incubated at 99 °C for 10 min. Protein digestion, peptide labelling, and MS-based proteomics were performed as previously described^90^. Limma analysis was performed similarly as previously described^90^ to determine proteins that were significantly up or downregulated.

### Orthology analysis of resistant strains

All complete genomes belonging to the *A. baumannii*, *E. coli* and *K. pneumoniae* species were downloaded from NCBI RefSeq using ncbi-genome-download^91^. Newly sequenced genomes were annotated using prokka, with default parameters^82^. The pangenome for each of the three species was computed separately using panaroo with the “--clean-mode strict -- merge_paralogs” options^92^. One or more protein sequences for each gene were then sampled in the three pangenomes, giving priority to parental sensitive strains. If a gene was not present in any of these “focal” strains, a random strain was selected. Sampled protein sequences were annotated using egnog-mapper, with the following parameters: “--target_orthologs one2one --go_evidence all --tax_scope Bacteria --pfam_realign realign”^93^. GO terms associated with each gene cluster were recovered using this automatic annotation. We further expanded this set by querying the NCBI protein database using Biopython’s Entrez interface^94^. This was possible because we used complete genomes from RefSeq. We then combined the two annotation sets to derive a set of GO terms for each gene cluster in the three species.

## Data availability

Source data for all figures is available with this manuscript. The mass spectrometry proteomics data have been deposited to the ProteomeXchange Consortium via the PRIDE partner repository with the dataset identifiers PXD050778 (reviewer id; OlbUlX9I) and PXD050827 (reviewer id; zckFx69i).

## Author contribution

E.C., M.T. and S.G. conceived and designed the study. S.G. and A.T. supervised the study. E.C., M.T., M.S. and S.G. designed the experiments. M.S. and S.G. performed and analysed the MIC testing of clinical isolates; M.T., M.S. and J.J.P. analysed resistant mutants. M.T. and J.P. performed time-kill experiments and *S.* Typhi intracellular killing assay. M.T. and M.S. performed efflux-pump inhibitor experiments. M.S. performed nitroxoline resistance evolution and sequencing of resistant clones. M.T. performed *G. mellonella* infection experiments. M.T., M.E., C.R., P.C. and S.G. performed the XRF experiment. E.C. performed and analysed the time-lapse microscopy, drug pretesting and combination screen with the help of M.K., *E. coli lptD4213* MIC testing, experiments on colistin-resistant *Enterobacteriaceae*, EOP and NPN assays. E.C. and A.O. performed the phylogeny analysis. A.M. performed the 2D-TPP and proteomics experiments and E.C. analysed the data. M.G. performed the orthology analysis of resistant strains. A.K. and F.C. performed the chemical genetic pretesting and screen; V.V. analysed the data. T.G.S. analysed the genome sequences and performed the SNP analysis. E.C. and S.G. wrote the manuscript with input from all authors. All authors approved the final version.

## Acknowledgements

S.G. was supported by the Rolf. M. Schwiete-Stiftung. A.T., V.V., F.C., A.B.-N. and M.Z. were supported by EMBL. M.K. was supported by Vetenskapsrådet 2019-00666. A.M. was supported by a fellowship from the EMBL Interdisciplinary Postdoc (EI3POD) programme under Marie Skłodowska-Curie Actions COFUND (grant number 664726). M.G. was funded by the Deutsche Forschungsgemeinschaft (DFG, German Research Foundation) under Germany’s Excellence Strategy - EXC 2155 - project number 390874280. A. O. and P.B. were supported by the German Federal Ministry of Education and Research (LAMarCK, 031L0181A to P.B.). M.M.S. was supported by the Allen Distinguished Investigator award through the Paul G. Allen Frontiers Group”. We thank ESRF for granting beamtime on ID16A through experiment LS-3269 (DOI 10.15151/ESRF-ES-1346211120).

## Extended Data Figures

**Extended Data Figure 1.**
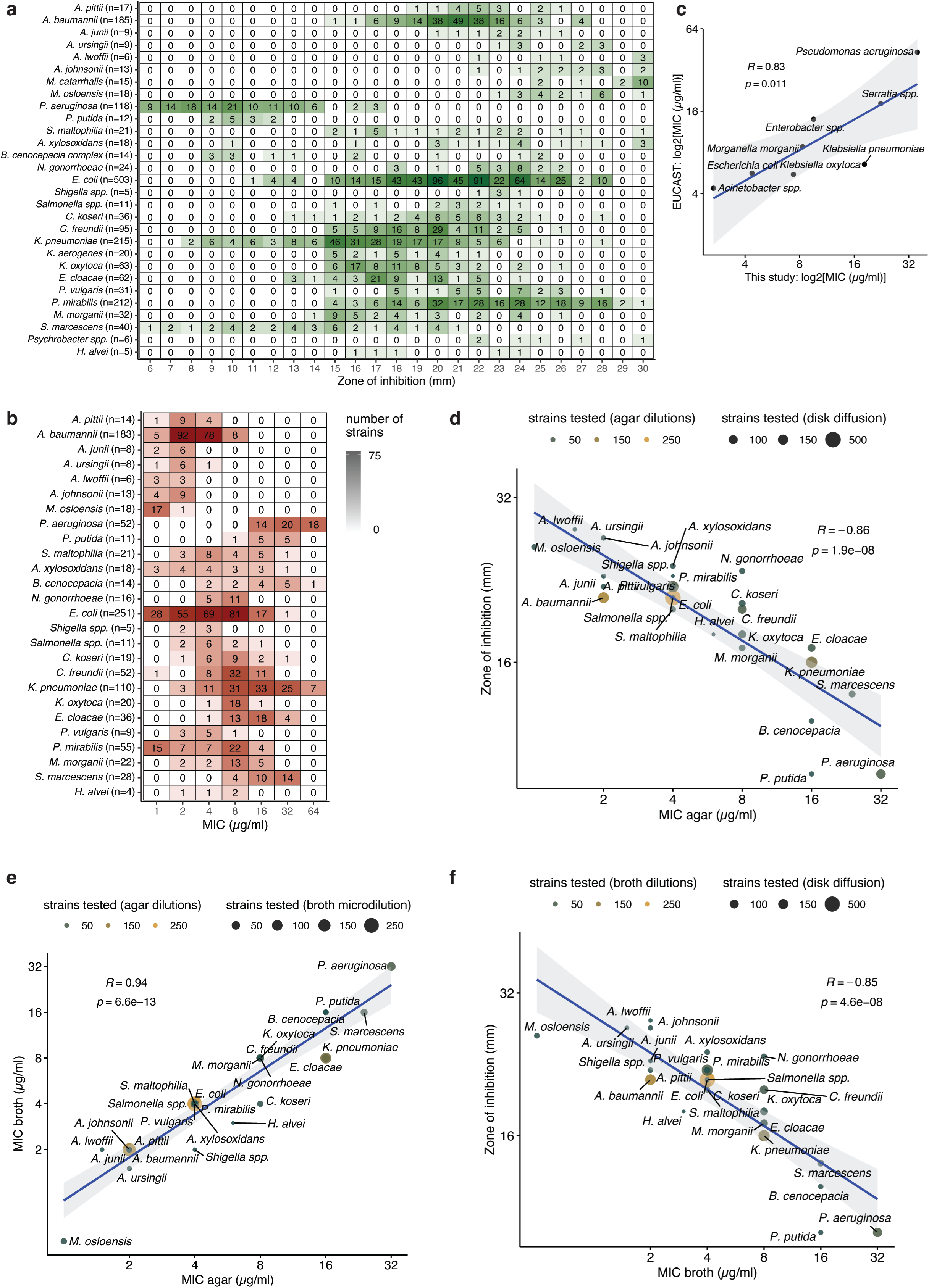
Nitroxoline MIC measurement via disk diffusion and agar dilutions and agreement across the different assays and with EUCAST. **a-b.** MIC measurements via disk diffusion (**a**) and agar dilution (**b**). Sensitivity is expressed as inhibition zone diameter (mm) (**a**) or µg/ml (**b**) (Methods). Each value is annotated and color-coded according to the number of strains it was measured in. Multiple species are aggregated for *Salmonella* spp., *Shigella* spp., *Burkholderia cenocepacia complex* and *Psychrobacter spp.* (Source Data ED Fig. 1a-b). **c.** Concordance between MICs measured in broth dilution in this study and according to EUCAST^19^ for the eight overlapping species. **d-f.** Concordance between MICs measured across three orthogonal methods: agar dilution and disk diffusion (**d**), broth and agar dilution (**e**), broth dilution and disk diffusion (**f**). Pearson’s correlation coefficient and p-value are shown. Disk diffusion results are expressed as in Extended Data Fig. 1a.

**Extended Data Figure 2.**
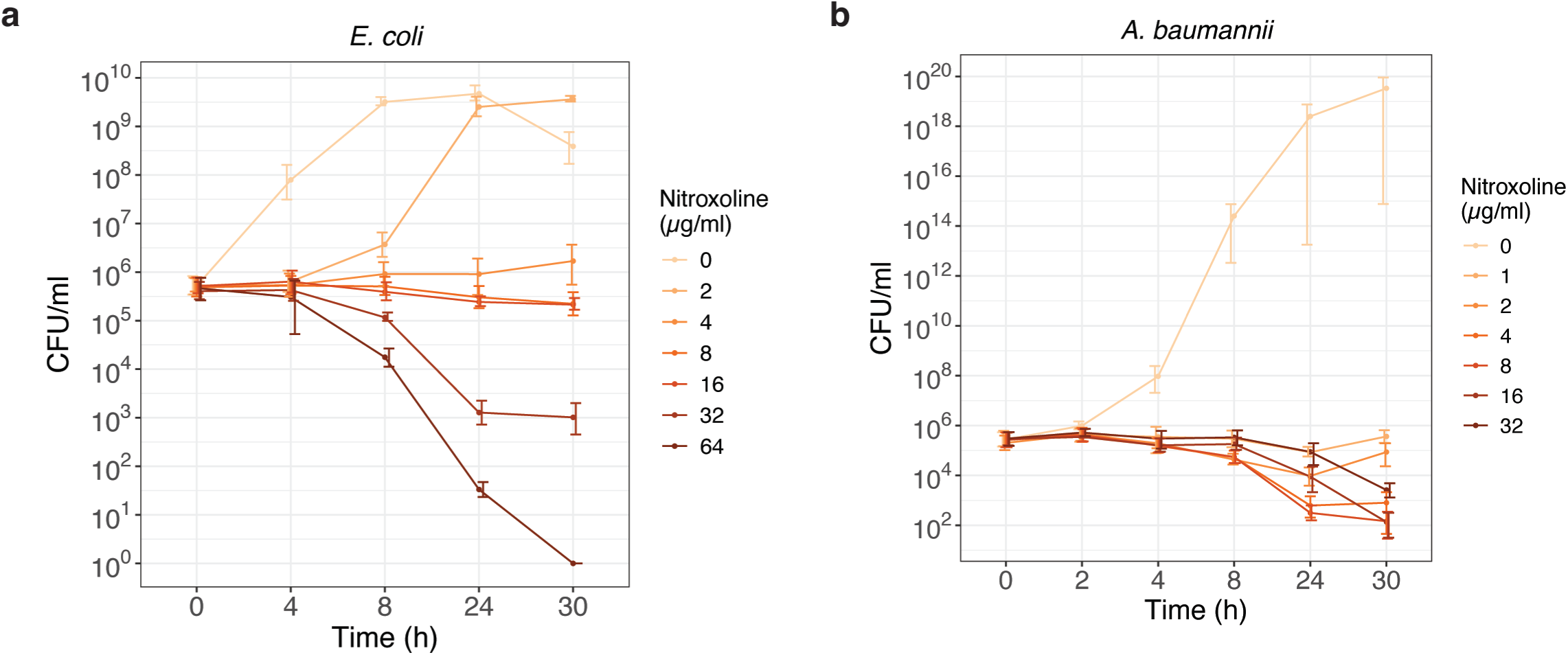
Bactericidal activity of nitroxoline. **a.** Nitroxoline is bacteriostatic against *E. coli* BW25113 at concentrations up to 4 x MIC (16 µg/ml) and bactericidal from 8 x MIC (32 µg/ml). Results are represented as in Fig. 1e. **b.** Time-kill curves for nitroxoline in *A. baumannii* ATCC 19606^T^ (Fig. 1e) including the no-drug control. Results are represented as in Fig. 1e.

**Extended Data Figure 3.**
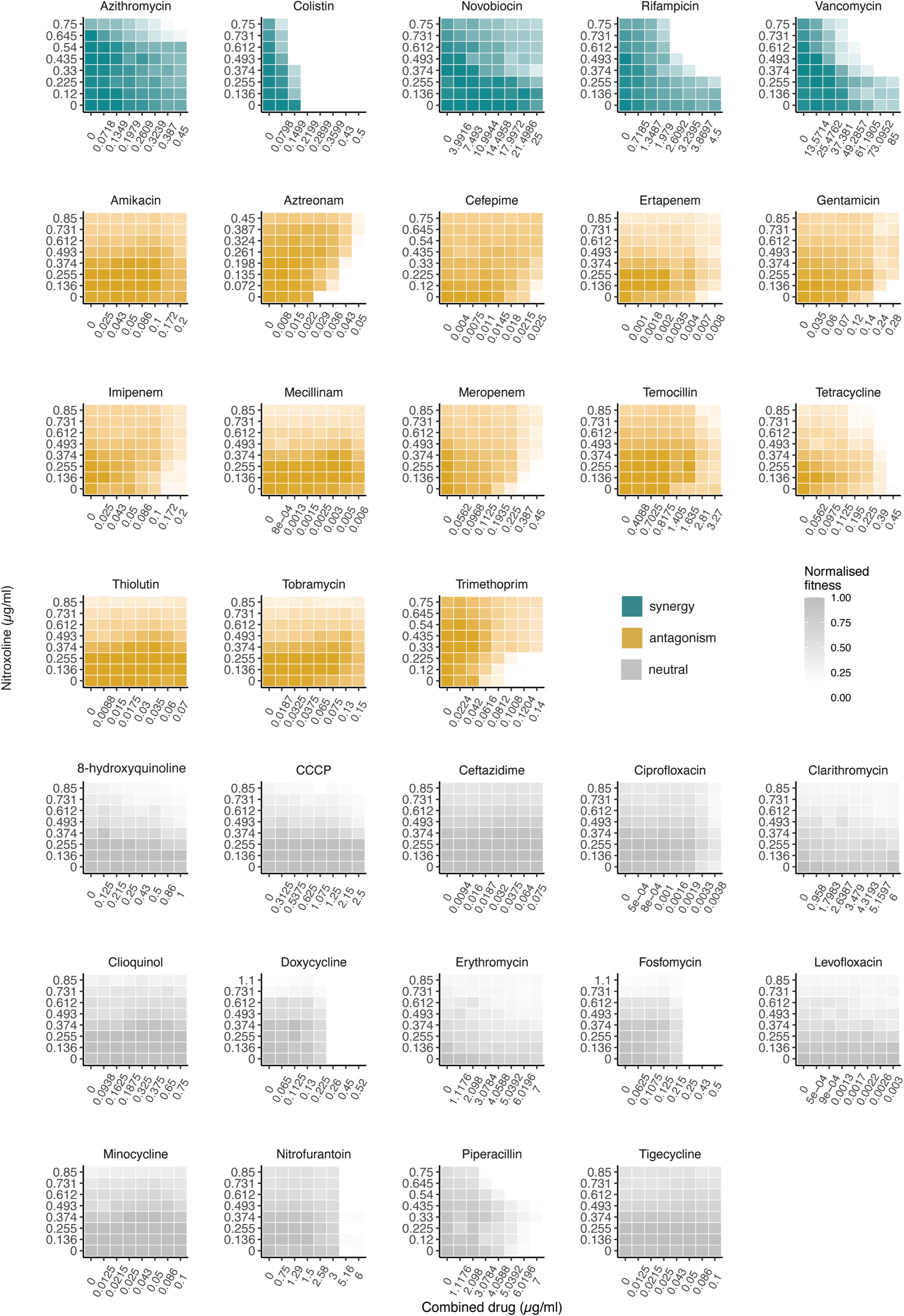
Checkerboard microdilution of nitroxoline combinations with 32 antimicrobials. Drugs were tested in combination with nitroxoline in 8 x 8 broth microdilution checkerboards. The median fitness (OD_595_ at 7.5 h normalized by no-drug controls) across at least two biological replicates is shown (**Supplementary File**).

**Extended Data Figure 4.**
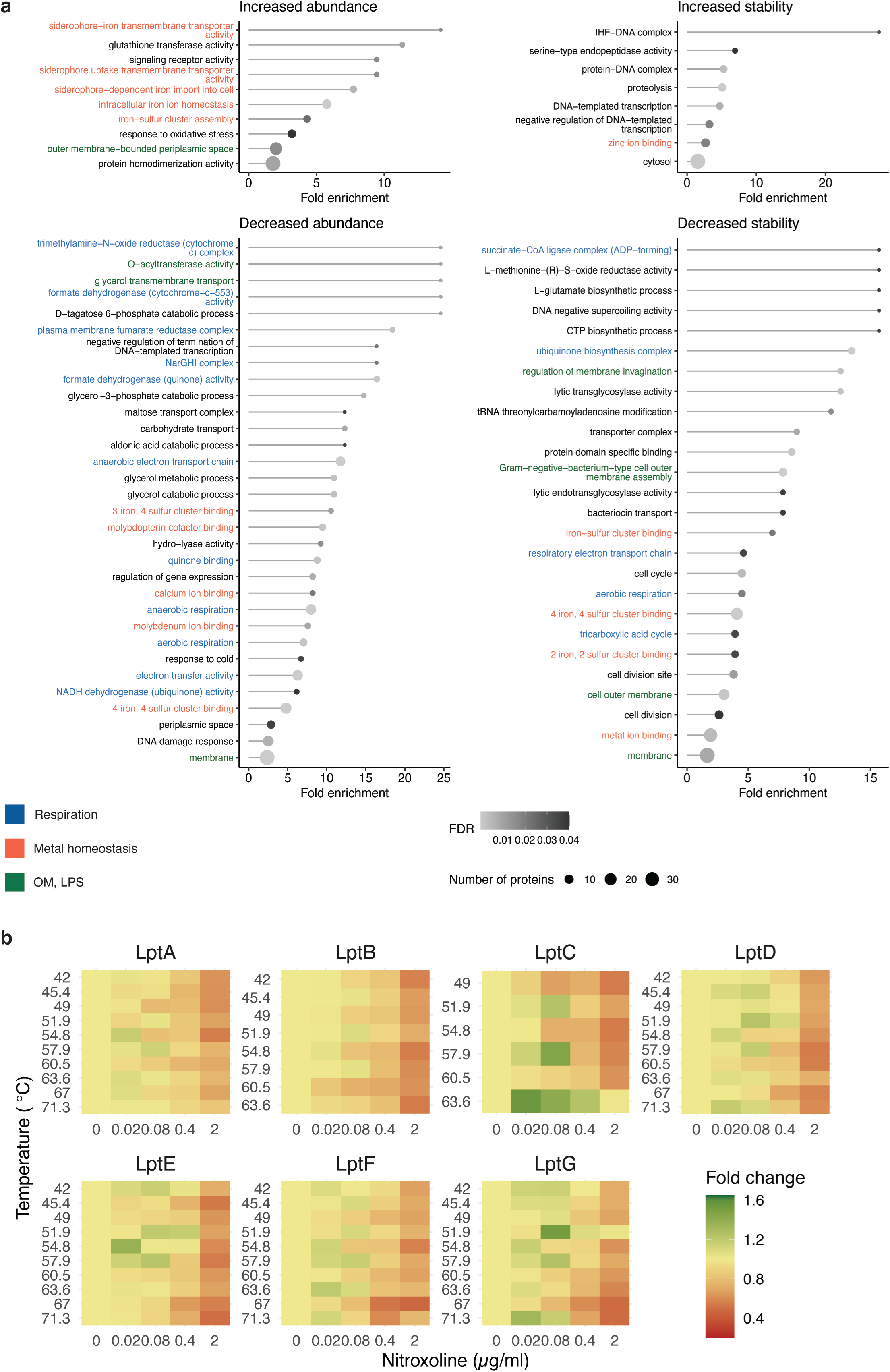
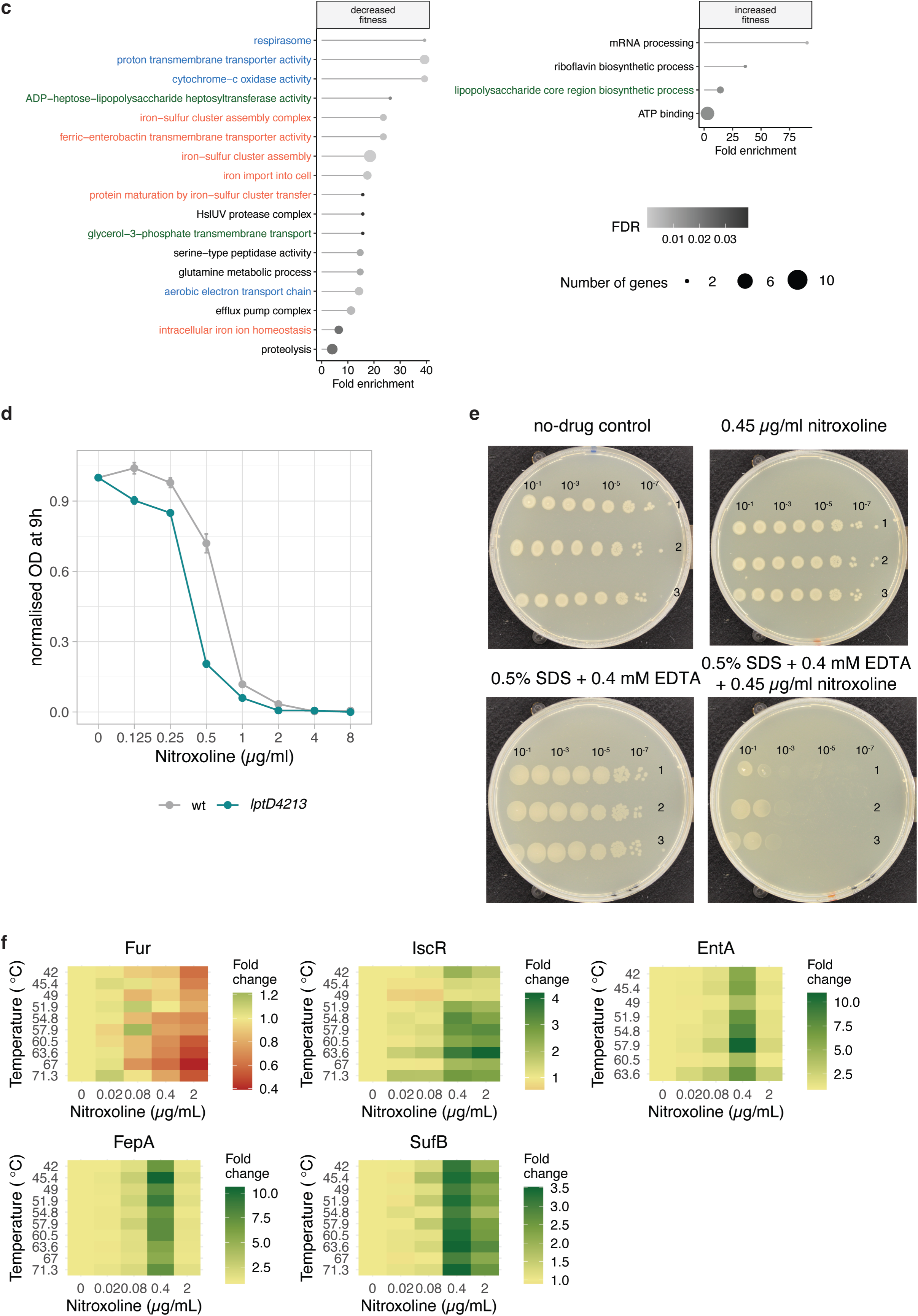
Nitroxoline perturbs the OM and affects metal homeostasis in *E. coli*. **a.** GO enrichment of 2D-TPP abundance and stability changes (Fig. 3a, **Supplementary Table 3**). Significantly enriched GO terms are shown (adjusted p-value < 0.05, Fisher’s exact test). Dot size represents the number of hit proteins for each term, dot colour the adjusted p-value, label colour highlights GO terms related to OM or LPS, metal homeostasis, and respiration. **b.** Nitroxoline affects the abundance and stability of the LPS transport machinery. In the thermal stability profiles of Lpt system members, protein fold change is shown for each temperature and nitroxoline concentration. **c.** GO enrichment of chemical genetics hits (Fig. 3b, **Supplementary Table 4**). Results are represented as in Extended Data Fig. 4a. **d.** Nitroxoline is more potent upon genetic perturbation of the OM. Nitroxoline susceptibility in *E. coli* BW25113 and its OM-defective derivative strain, carrying the *lptD4213* mutation^40^. Data is shown as in Fig. 2b. For full growth curves see Supplementary File. **e.** Lower concentrations of nitroxoline and EDTA were tested than in Fig. 3d (EOP assay, Methods) to show the first dosage at which growth was visible upon combination. **f.** Nitroxoline induces effects in 2D-TPP consistent with iron-sulfur cluster disruption. Thermal profiles are represented as in Extended Data Fig. 4b.

**Extended Data Figure 5.**
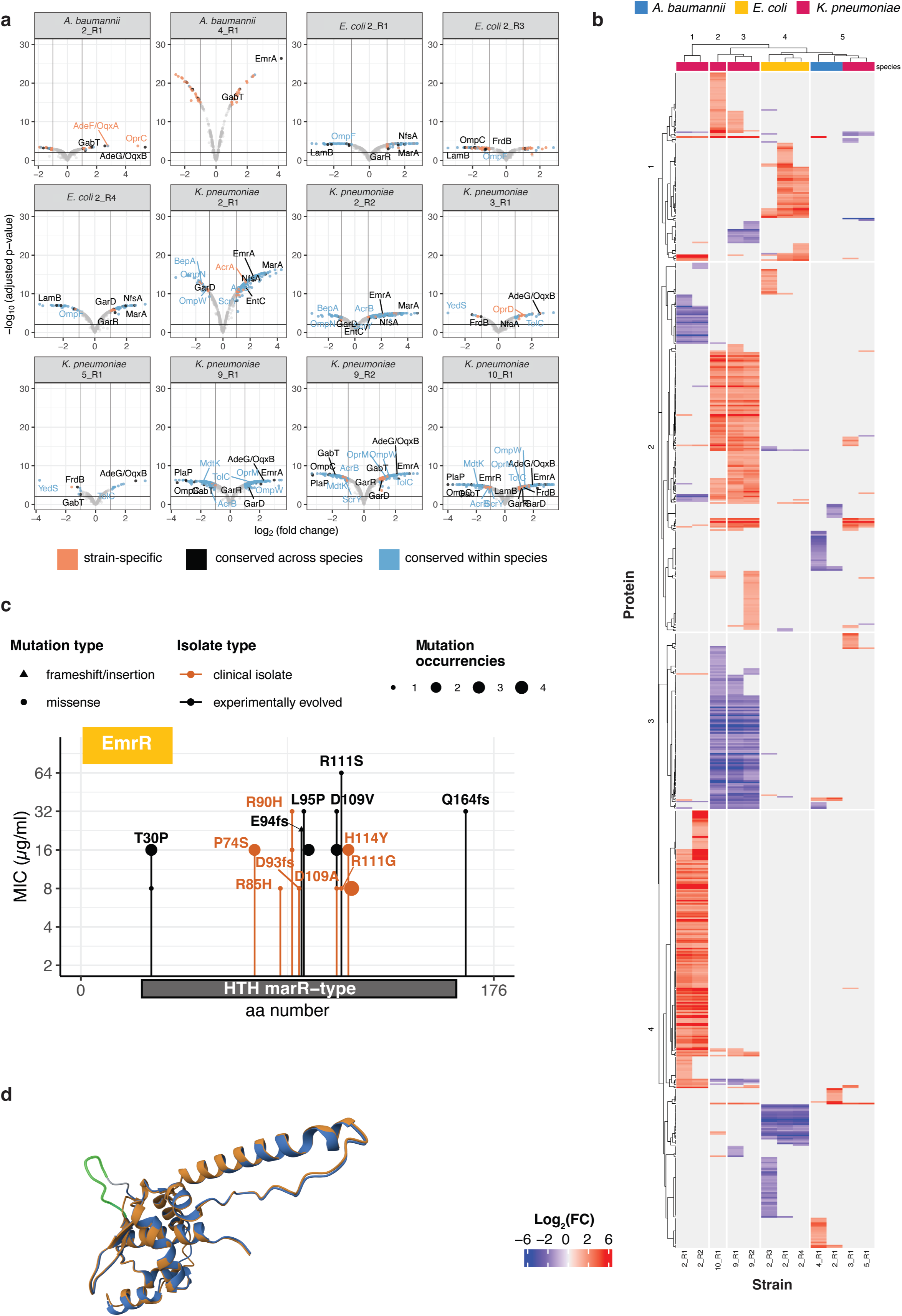
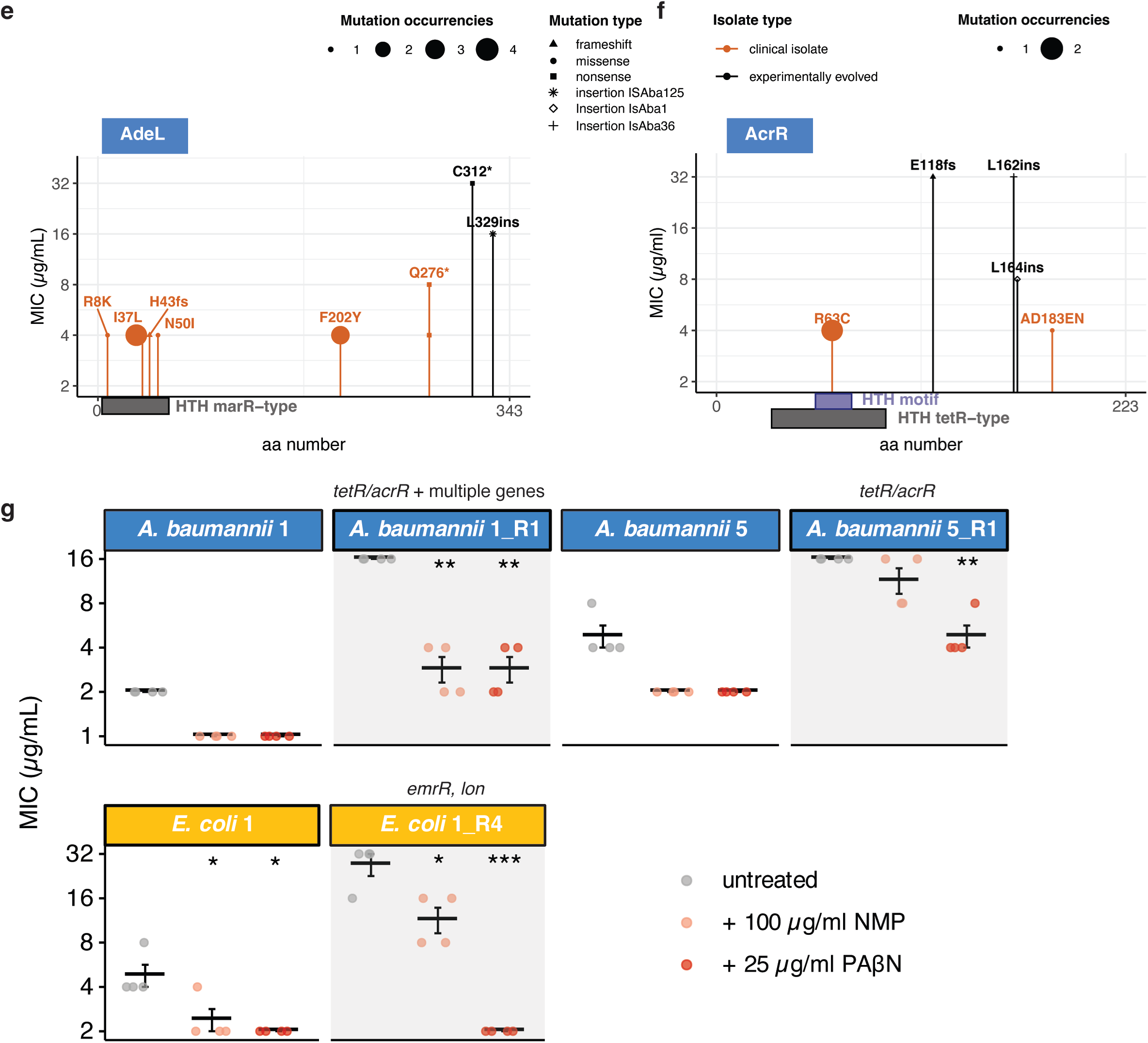
Nitroxoline resistance is based on conserved mechanisms across species. **a.** Volcano plots of abundance changes in proteomics of resistant strains compared to isogenic sensitive strains. Effect size and statistical significance (Methods) are represented on the x- and y- axis, respectively. Data points are colour-coded according to their conservation across or within-species. For sensitive strains see Supplementary File. **b.** Protein abundance changes in nitroxoline-resistant strains. Only significant and strong changes (adjusted p-value ≤ 0.01, |log_2_ fold-change (abundance)| ≥ 2) are included (**Supplementary Table 5**). Proteins and strains are clustered according to Pearson’s correlation. Results are represented as in Fig. 5b. **c.** Amino acid changes from *emrR* mutations in clinical and experimentally evolved nitroxoline-resistant isolates. Results are represented as in Fig. 5d. **d.** Superimposition of mutated and wild-type OqxR structure. The G60-L67 duplication loop is highlighted in green (**Methods**, **Supplementary File**). **e-f.** Amino acid changes from *adeL* (**e**) and *acrR/tetR* (**f**) mutations in clinical and experimentally evolved nitroxoline-resistant isolates. Results are represented as in Fig. 5d. **g.** Efflux-pump inhibitors resensitize nitroxoline-resistant strains. Results are represented as in Fig. 5e.

## Supplementary Tables

**Supplementary Table 1.** All the strains used in this study, their genotype, antibiotic resistance and nitroxoline MIC, except for those shown in Fig. 1b-c and Extended Data Fig. 1a-b (see Source Data).

**Supplementary Table 2.** The drugs tested in combination with nitroxoline in *E. coli* BW25113, their chemical class and highest concentration tested.

**Supplementary Table 3.** 2D-TPP data from *E. coli* BW25113 whole-cell samples exposed to nitroxoline (Methods).

**Supplementary Table 4.** Mutant fitness as log(fold change) and FDR from the chemical genetic screen performed on the *E. coli* Keio collection with nitroxoline (Methods).

**Supplementary Table 5.** Proteomics data from nitroxoline-sensitive and resistant strains listed in Supplementary Table 1 and annotated with their orthologous groups (Methods).

## Supplementary File

Supplementary Figs. 1-9.

## Supplementary Videos

**Supplementary Videos 1-2.** Time-lapse of *A. baumannii* growing on cation-adjusted Mueller-Hinton-agarose 1% pad, supplemented (**1**) or not (**2**) with 8 µg/ml nitroxoline.

