## Supplementary File for "Uncovering nitroxoline activity spectrum, mode of action and resistance across Gram-negative bacteria"

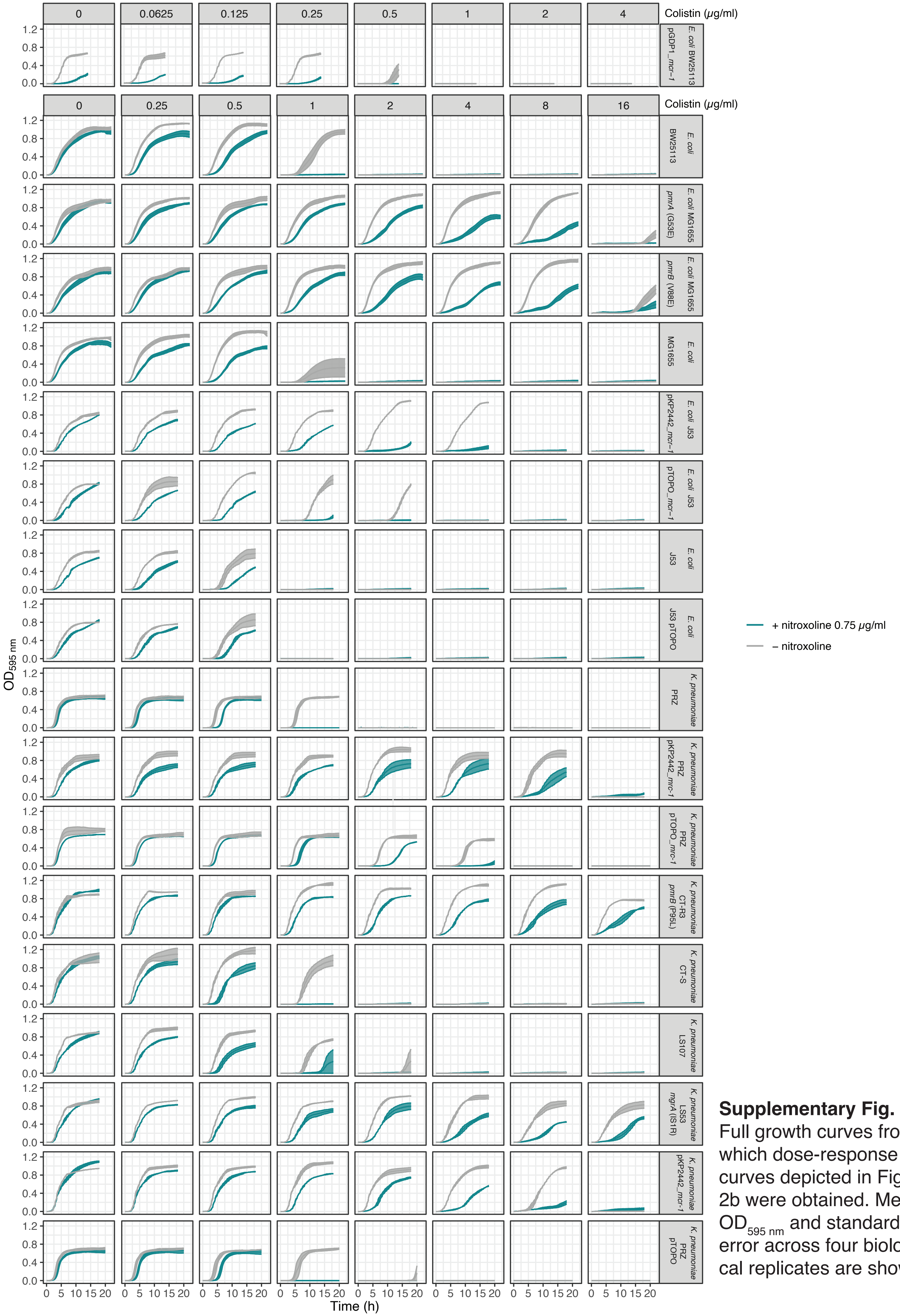


**Supplementary Fig. 2.** Full growth curves from which dose-response curves depicted in Fig. 2b were obtained. Mean $OD_{595\ nm}$ and standard error across four biological replicates are shown.

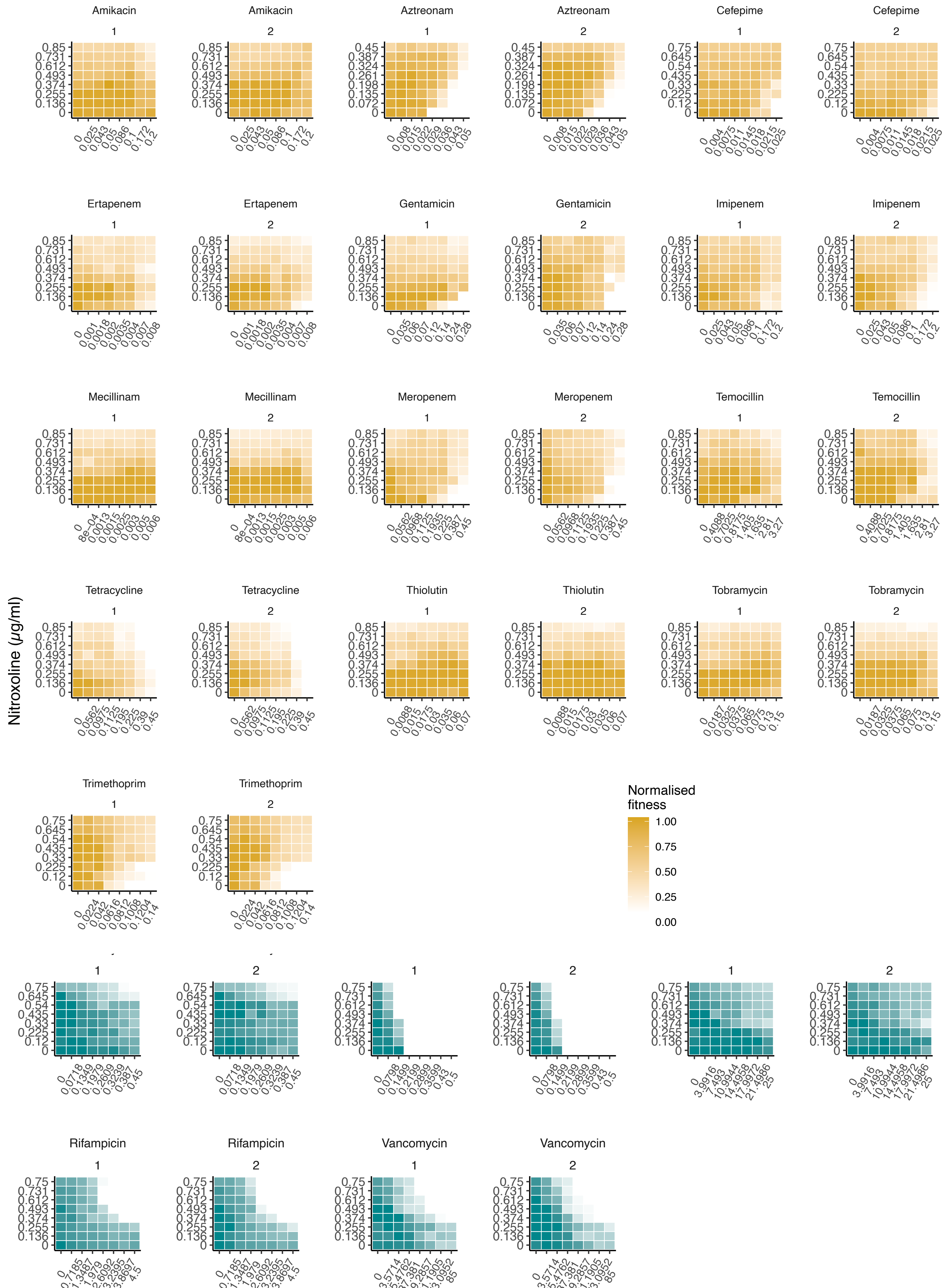


**Supplementary Fig. 1**

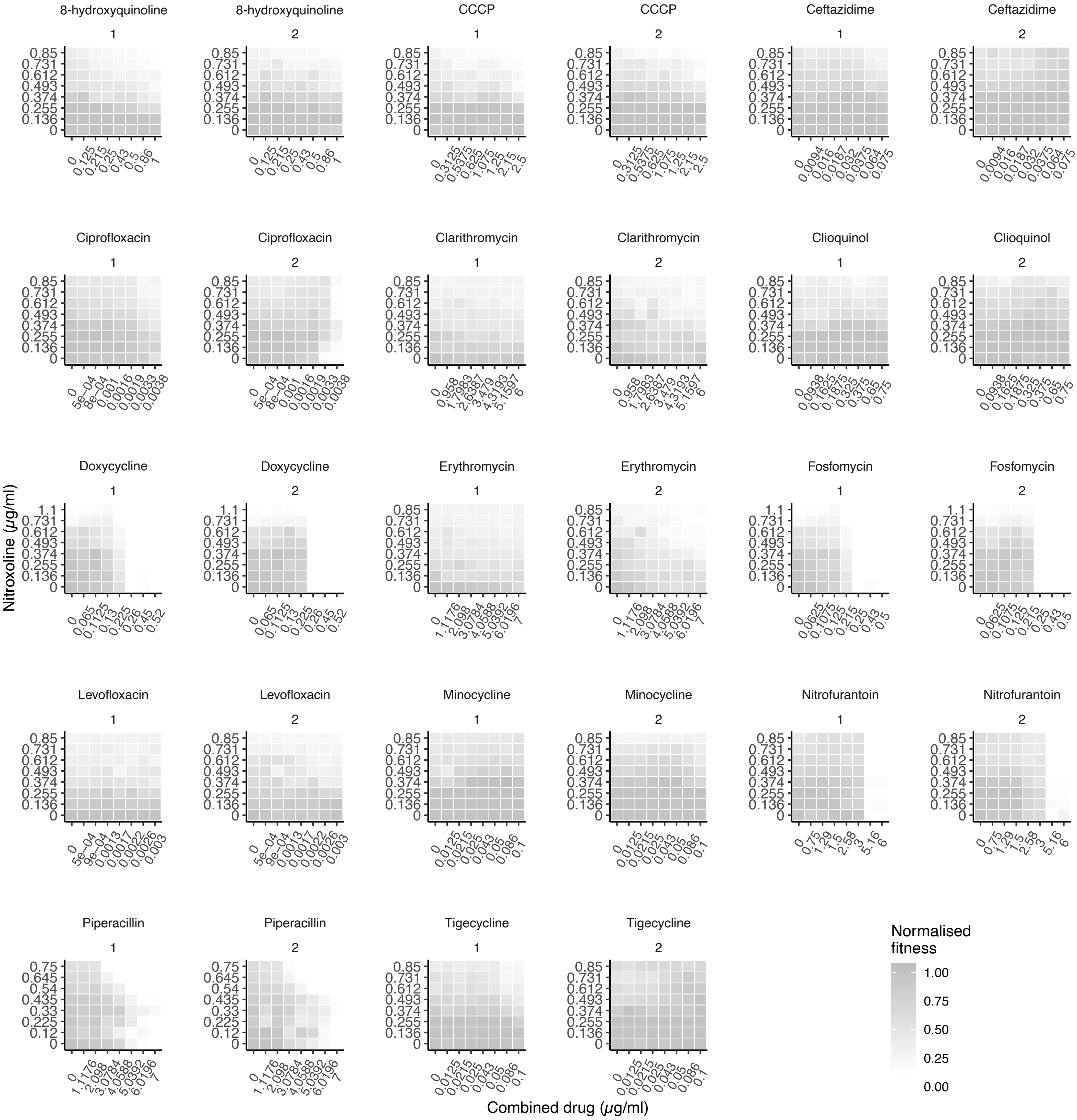


**Supplementary Fig. 1 - continued.** Two biological replicates of checkerboard assays displayed in Extended Data Fig. 3, from which Bliss interaction scores shown in Fig. 2a are obtained. Results are obtained and represented as in Extended Data Fig. 3.

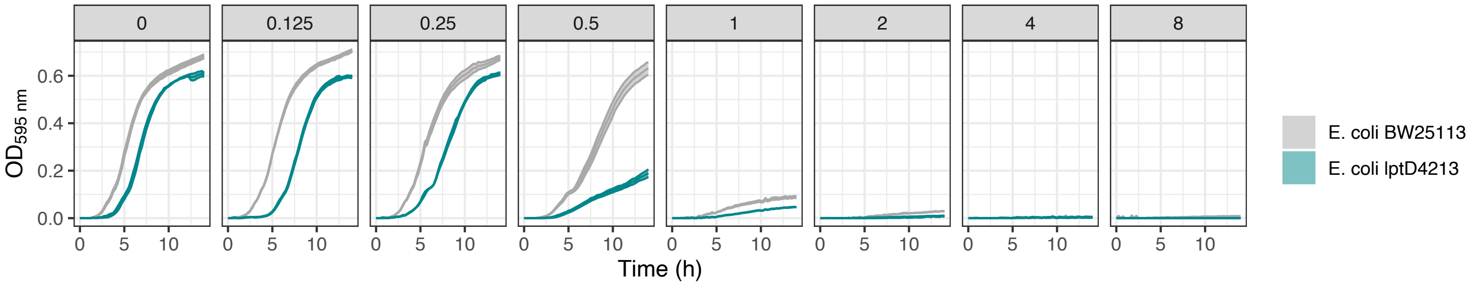


**Supplementary Fig. 3.** Full growth curves from which dose-response curves depicted in Extended Data Fig. 4d were obtained. Mean $OD_{595\ nm}$ and standard error across four biological replicates are shown.

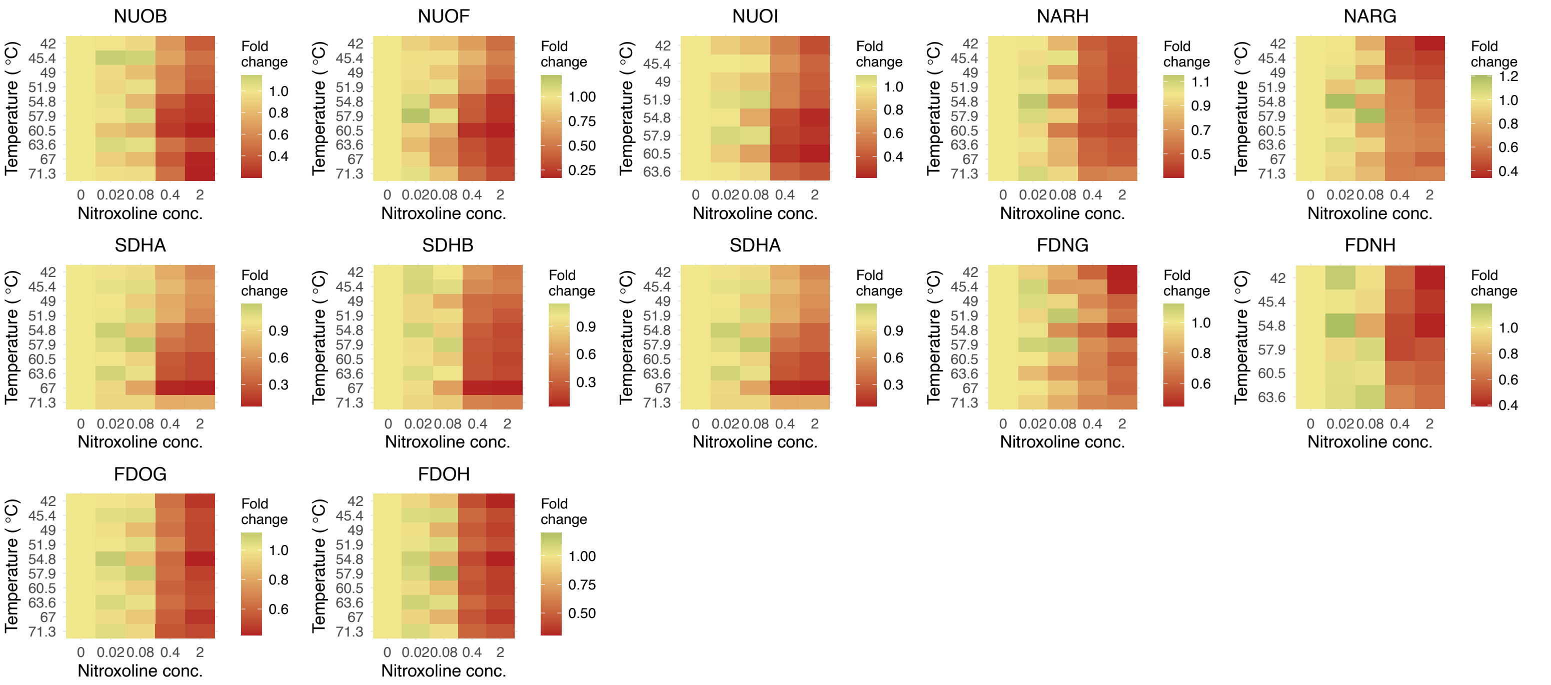


**Supplementary Fig. 4.** Thermal stability protiles of members of the respiratory chain. Data is represented as in Fig. 4b.

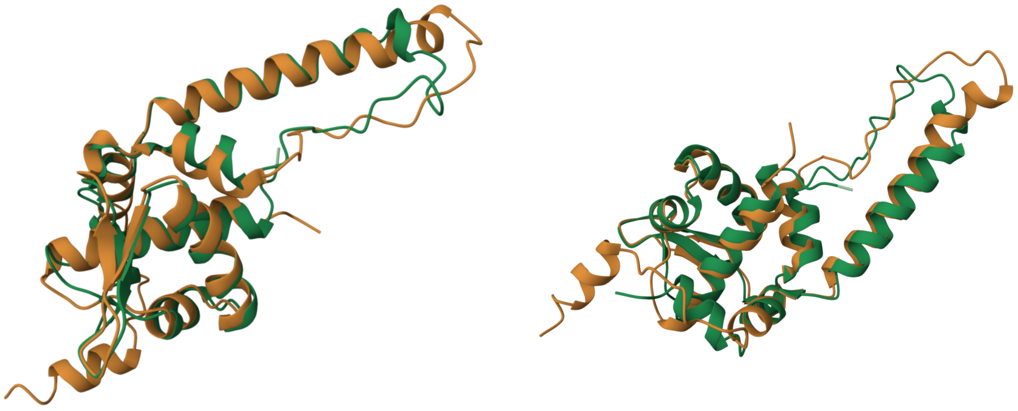

**Supplementary Fig. 5.** Structural alignments between OqxR and NsrR (Methods), which was used for the domain annotation shown in Fig. 5d.

**wild-type OqxR:**
MLDYRFPTALQMVLSVAMAEQMGERSTSAILAYGLEANPSFIRKLMVPLTRDGIIVSTLGRNGSIHLGRPADKITLRDIYLSVIEDKKL
WASRPDVPARCVVSANACWYFKSVADEAEQASLNVLARHTAASALEAVKNADTSGCDPVPEMIARFKKAH

**OqxR G60-L67dup:**
MLDYRFPTALQMVLSVAMAEQMGERSTSAILAYGLEANPSFIRKLMVPLTRDGIIVSTLGRNGSIHLGRNGSIHLGRPADKITLRDIY
LSVIEDKKLWASRPDVPARCVVSANACWYFKSVADEAEQASLNVLARHTAASALEAVKNADTSGCDPVPEMIARFKKAH

**Supplementary Fig. 6.** Amino acid sequences of wild-type OqxR and OqxR carrying the G60-L67 duplication (red), depicted in Extended Data Fig. 5d.

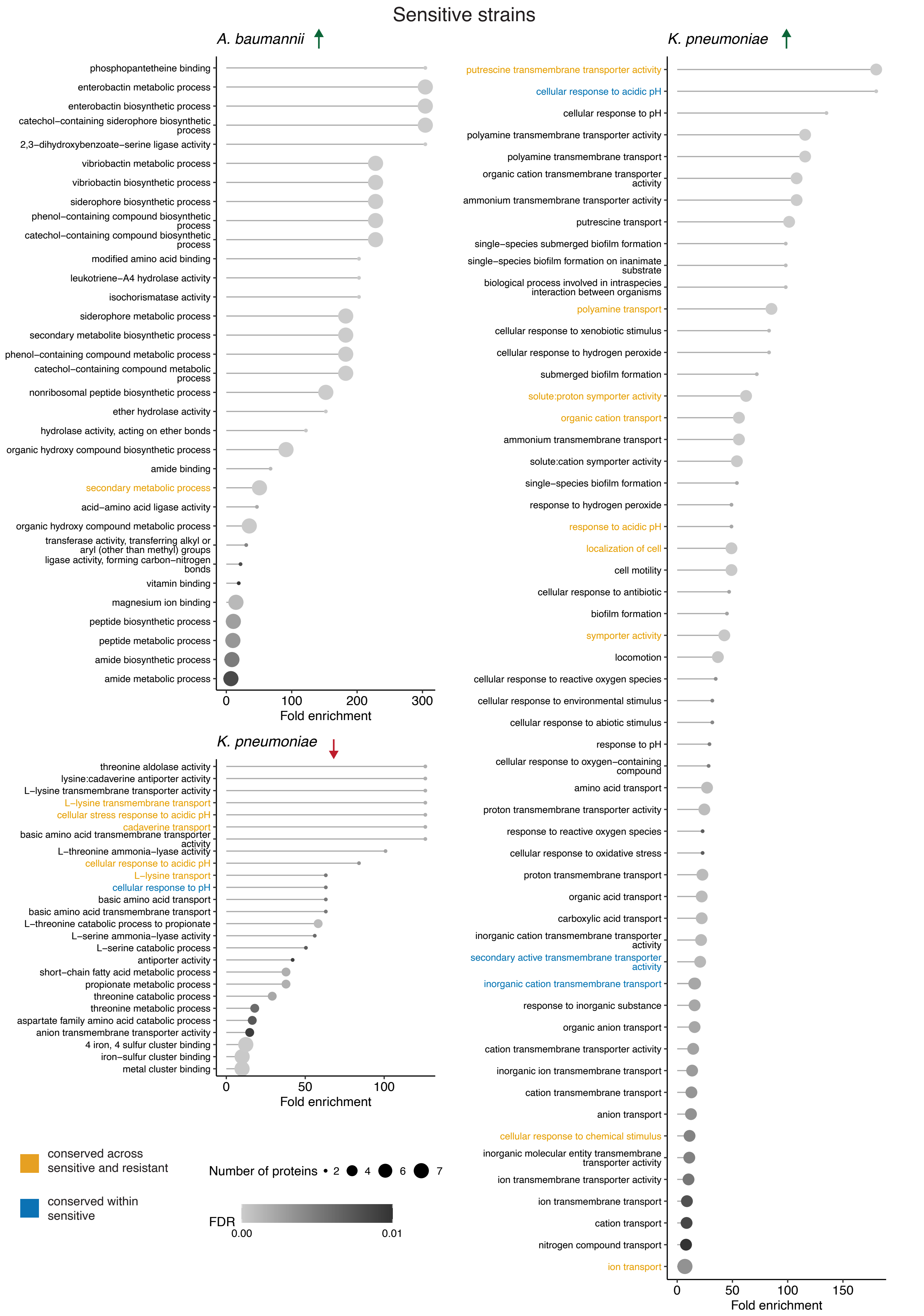


**Supplementary Fig. 8**. GO enrichment of significant hits from proteomics on nitroxoline-sensitive strains. Data is represented as in Supplementary Fig. 7.

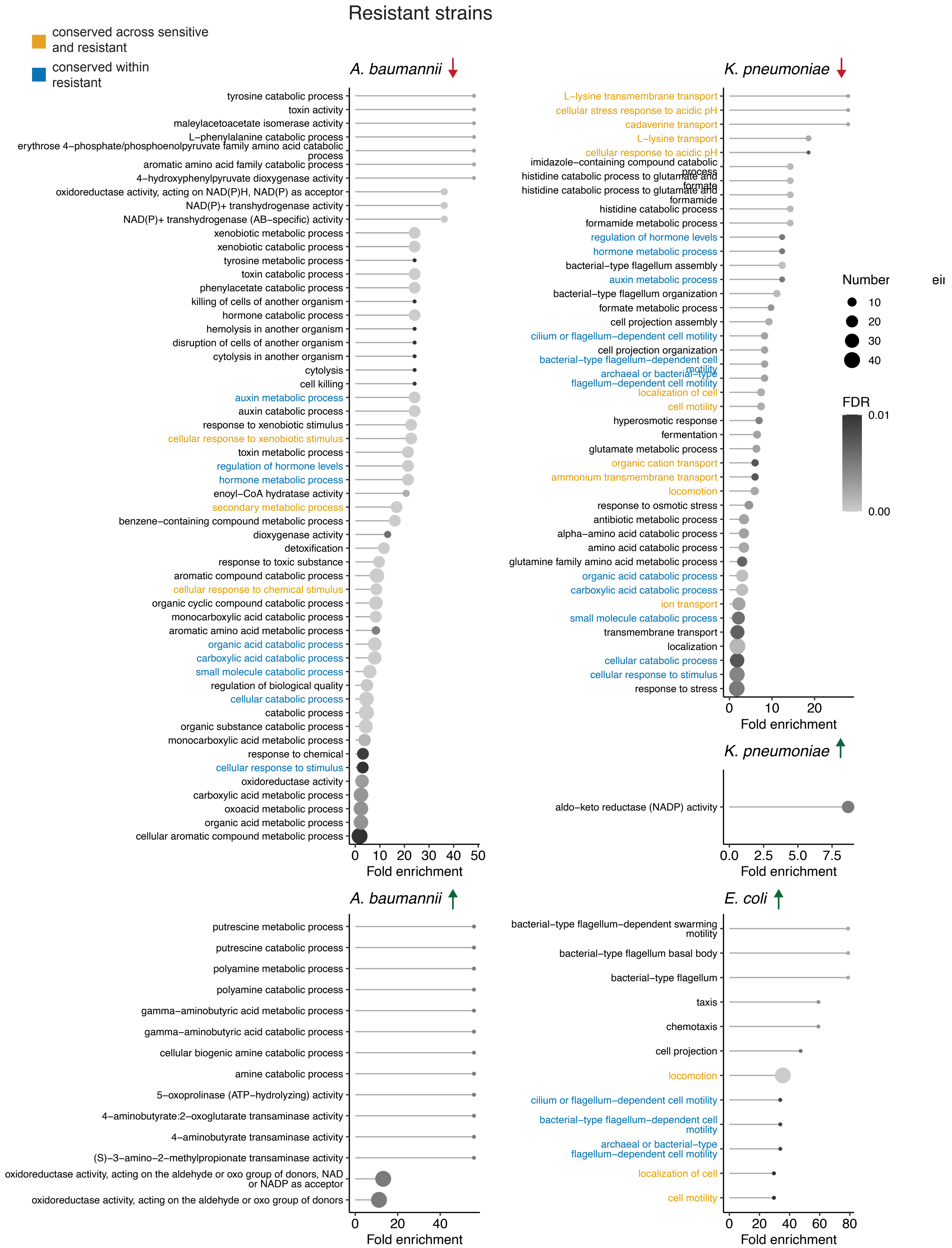


**Supplementary Fig. 7**. GO enrichment of significant hits from proteomics on nitroxoline-resistant strains (Fig. 5b, Extended Data Fig. 5b, Supplementary Table 5). Only sets yielding significant enrichments (down- or up-regulation in the indicated species) are shown (adjusted p-value $< 0.05$, one-sided Fisher's exact test). The number of protein hits is annotated for each term.

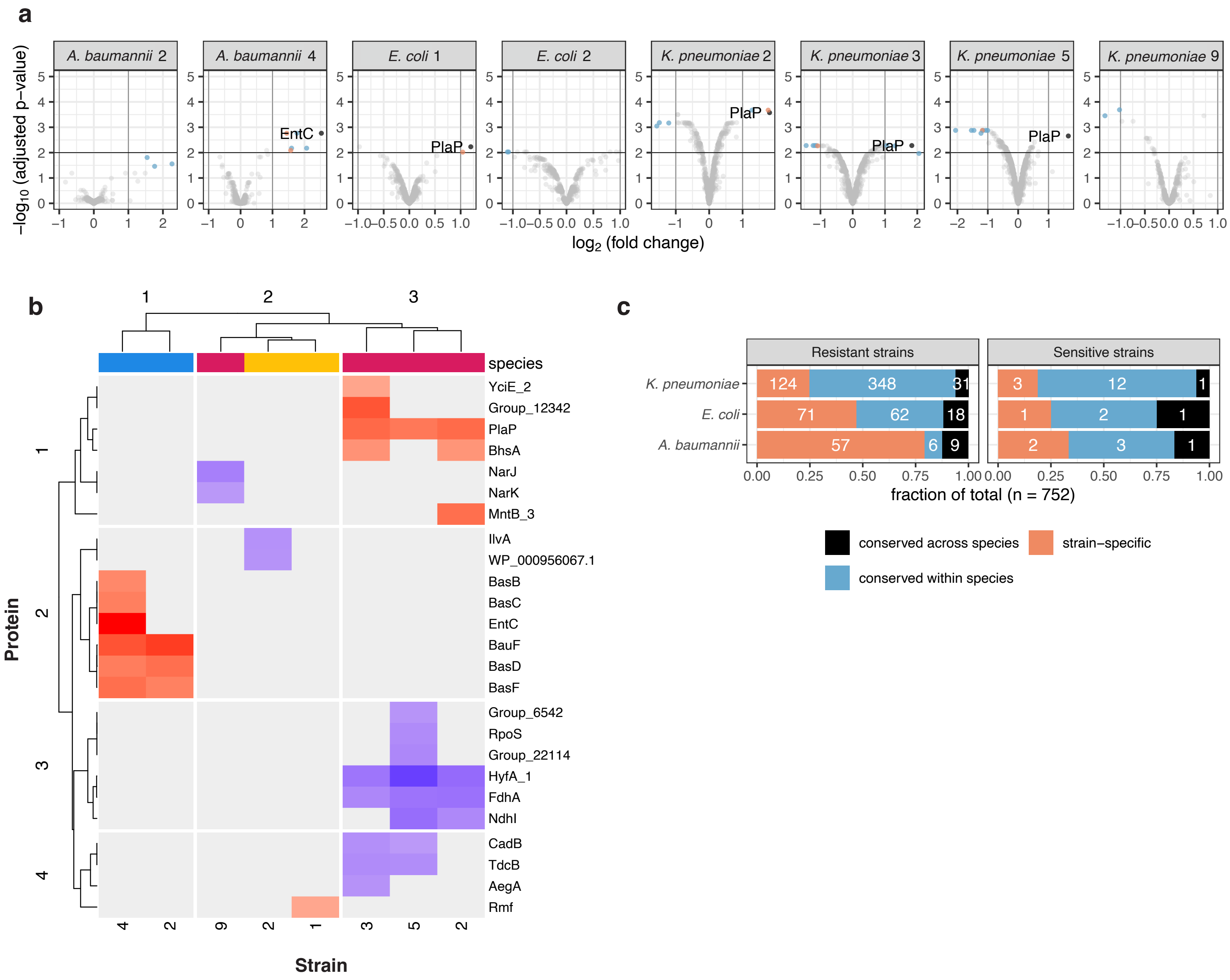


**Supplementary Fig. 9. a.** Volcano plots of abundance changes in proteomics of sensitive strains exposed to nitroxoline compared to untreated controls. Data is represented as in Extended Data Fig. 5a. **b.** Protein abundance changes in nitroxoline-sensitive strains. Data are represented as in Extended Data Fig. 5b. **c.** Conservation of protein changes in resistant and isogenic sensitive strains exposed to nitroxoline, color-coded as in Extended Data Fig. 5a.
